# Ion and water permeation through Claudin-10b paracellular channels

**DOI:** 10.1101/2024.07.03.601692

**Authors:** Alessandro Berselli, Giulio Alberini, Fabio Benfenati, Luca Maragliano

## Abstract

The structural scaffold of epithelial and endothelial tight junctions (TJ) comprises multimeric strands of claudin (Cldn) proteins, which anchor adjacent cells and control the paracellular flux of water and solutes. Based on the permeability properties they confer to the TJs, Cldns are classified as channel- or barrier-forming. Some of them, however, show mixed features. For instance, Cldn10b, expressed in kidneys, lungs, and other tissues, displays high permeability for cations and low permeability for water. Along with its high sequence similarity to the cation- and water-permeable Cldn15, this makes Cldn10b a valuable test case for investigating the molecular determinants of paracellular transport. In lack of high-resolution experimental information on TJ architectures, here we use Molecular Dynamics simulations to study two atomistic models of Cldn10b strands and compare their ion and water transport with those of Cldn15. Our data, based on extensive standard simulations and Free Energy calculations, reveal that both Cldn10b models form cation-permeable pores narrower than Cldn15, which, together with the stable coordination of Na^+^ ions to acidic pore-lining residues (E153, D36, D56), limit the passage of water molecules. By providing a mechanism driving a peculiar case of paracellular transport, these results provide a structural basis for the specific permeability properties of Cldn isoforms that define their physiological role.

**Graphical abstract:** 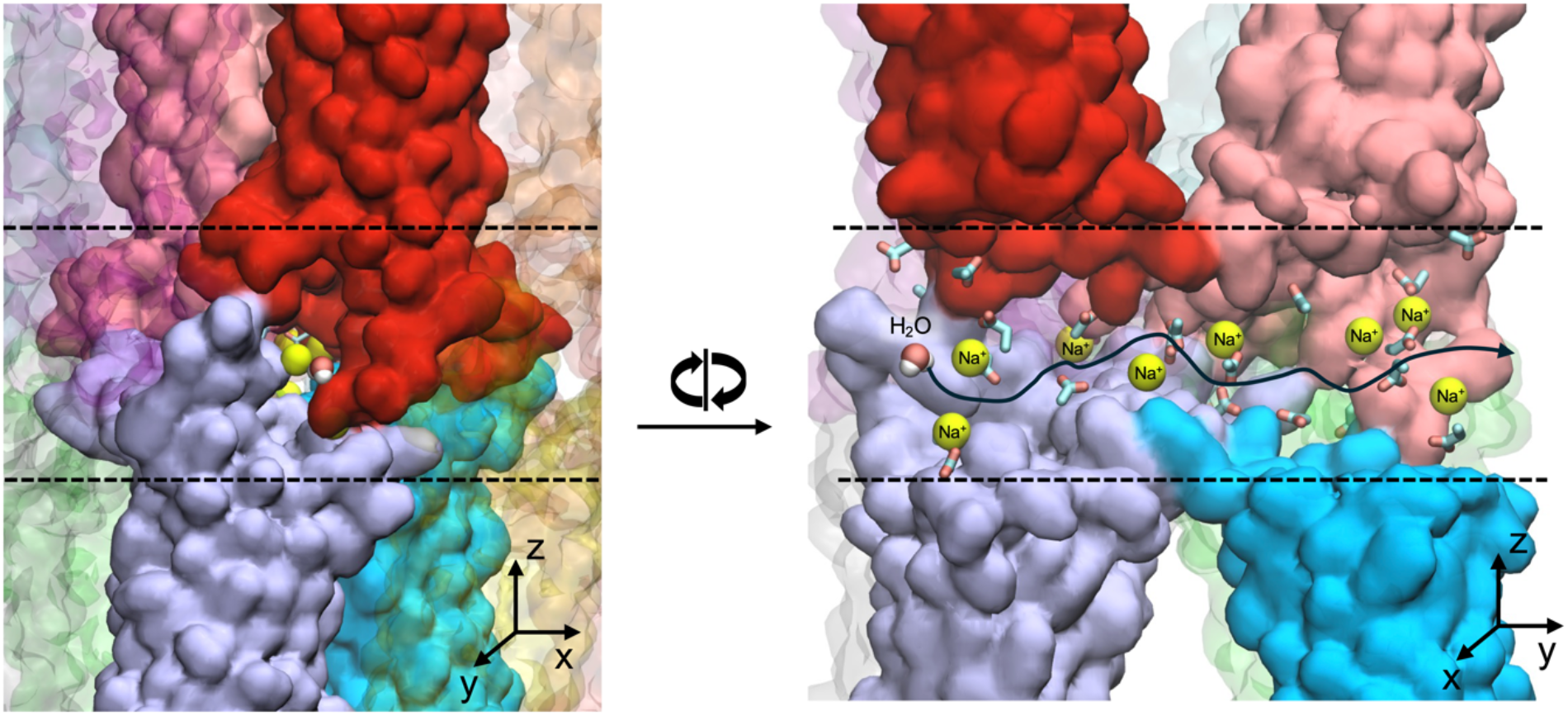

## Introduction

The movement of water, ions, and molecules across epithelia or endothelia occurs by following distinct transcellular and paracellular pathways. The former involves specific channels or transporters ^[1–4]^, while the latter relies on passive diffusion through the narrow space between adjacent cells, regulated by a meshwork of multiprotein strands hinging cells together and named tight junctions (TJs) ^[5–7]^. The TJ backbone is formed by multimers of claudin (Cldn) proteins that determine the selective permeability of the strands to electrolytes and molecules in a tissue-specific manner ^[8–10]^. Structurally, Cldns are folded in a transmembrane four-helix bundle (TM1-4), linked to each other by two loops in the paracellular space (ECL1-2) and one intracellular loop (ICL) in the cytosol, where the N- and C-termini are also found ^[11–14]^. The ECL1 is typically composed of ∼ 50 amino acids and comprises a four-stranded *β*-sheet (*β*1-4), along with a highly conserved extracellular helix (ECH). This domain is crucial for the formation of the intercellular Cldn-Cldn *trans*-interactions and includes the amino acids responsible for the TJ charge-selective permeability. On the other hand, the ECL2 is essential for the stabilization of the intracellular *cis*-interactions network ^[15,16]^. Shorter than ECL1 (< 30 amino acids), it provides a fifth *β*-strand (*β*5) to the extracellular five-stranded *β*-sheet. The family of mammalian Cldns includes 27 members ^[12,17]^, that can be classified based on their physiological function as channel-forming or barrier-forming. For example, Cldn2 ^[18]^, Cldn10b ^[19,20]^, Cldn15 ^[21]^, and Cldn21 ^[22]^ form cation-permeable systems, while Cldn4 ^[23]^, Cldn10a ^[24]^, Cldn17 ^[25]^ are selective to anions. Conversely, a barrier function was demonstrated for Cldn1 ^[26]^, Cldn3 ^[27]^, Cldn5 ^[28]^, Cldn11 ^[29]^ and Cldn14 ^[30]^. The evaluation of TJ water permeability is more controversial, given the difficulties in determining how the paracellular and transcellular routes contribute to the total transepithelial flux ^[31]^. Cldn2 and Cldn15 are thought to be water permeable ^[32,33]^, whereas Cldn17 ^[25]^, Cldn10a, and Cldn10b are not ^[31,33]^. The latter, largely expressed in the water-impermeable regions of the kidney Henle’s loop ^[34]^, is particularly intriguing since it shares a high sequence similarity with Cldn15.

While the structures of several Cldn monomers have been obtained experimentally, no detailed information on how they associate to form TJs is currently available. A model was suggested years ago ^[35]^, based on the Cldn15 monomeric crystal structure (PDB ID: 4P79 ^[36]^). The architecture displays two patterns of protein-protein interactions between monomers from the same cell, named *cis-linear* and *face-to-face* ^[37]^. The *cis-linear* arrangement generates single filaments of Cldn monomers, while *face-to-face* interactions join two antiparallel filaments in a double-row, where the ECL *β*-strands of each monomer couple form a distinctive “*half pipe*”-shaped structure ^[35]^. If double-row strands from neighbor cells *trans*-associate, the “*half pipes*” are joined, resulting in *β*-barrel pores parallel to the facing cell membranes. This architecture is referred to as the Suzuki, or joined-double row (JDR) ^[14]^ model, and is consistent with cross-linking experiments and freeze-fracture electron microscopy. It was refined and validated using Molecular Dynamics (MD) simulations first for Cldn15 ^[38–41]^, and then for several other isoforms ^[42–49]^. In our previous MD studies, we assessed it by calculating the free energy (FE) of ion permeation through single-pore Cldn tetramers, considering these as the minimal functional units of the model. Other groups analyzed higher-order multimers and reproduced ionic currents flowing through the pores ^[40,46]^. Altogether, these works showed that the model correctly reproduces the experimentally determined ion selectivity of various Cldns, which stems from the electrostatic environment generated by the pattern of charged pore-lining residues. In this study, we use all-atom MD simulations and FE calculations to determine whether the JDR model adequately captures the cation selectivity and low water permeability of Cldn10b. We employ two variants of a triple-pore multimer and compare results with those obtained for Cldn15. The two models differ by the orientation of the *β*1*β*2 loop (between the *β*1 and *β*2 strands of each monomer) that is not present in the original JDR structure, resulting in distinct orientations of critical pore-facing residues. Model1 is based on the Cldn15 double-pore conformation we previously published ^[39]^, while Model2 has been proposed by Piontek and collaborators ^[46,47]^.

Standard MD simulations on the microsecond timescale show that both Cldn10b JDR models are cation-selective and less water permeable than Cldn15, consistent with experimental findings. Water passage is hindered by the small size of the pores (∼ 2.2 Å of minimal radius) and the clustering of cations within the cavities. FE calculations result in repulsive barriers for Cl^-^ and attractive minima for cations, and barriers for water whose height depends on the number of Na^+^ ions occupying the pore. These outcomes support the validity of the JDR model, and provide insight into the structural and molecular determinants of the transport of ions and water by a distinctively selective Cldn, describing a novel mechanistic perspective of epithelial transport.

## Results

### Cldn-10b models are stable during MD simulations and exhibit different orientation of pore-lining residues

The two Cldn10b models, Model1 and Model2, embedded in the respective simulation box are shown in **Figure 1**. For each system, we ran three independent, 1-μs long, trajectories. We assessed the protein structural stability in the paracellular cleft by calculating the time evolution of the RMSD of pore-forming ECL domains. In **Figure 2a**,**b**, we report the average RMSD values for all ECL domains (upper panels) and the values for individual pores (lower panels). All profiles display a plateau, indicative of stable conformations.

**Figure 1.**
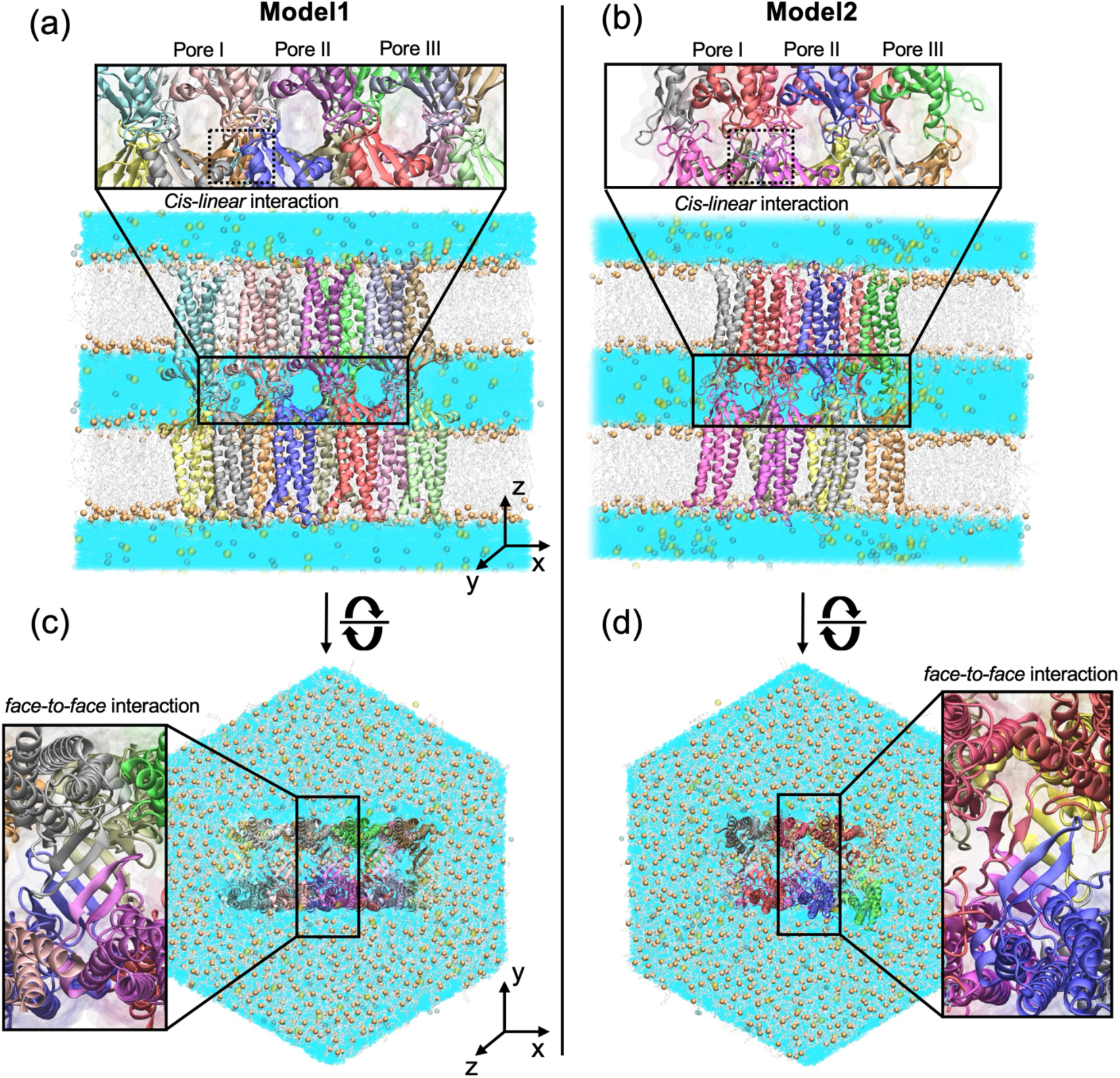
Claudin-10b triple-pore models. (**a**,**b)** Apical views of the Cldn10b triple-pore Model1 and Model2 structures, respectively. Close-up views of the ECL domains are shown, with the cis-linear interactions indicated with a dotted square. **(c**,**d)** Lateral views of Model1 and Model2, respectively. Close-up views of the face-to-face interaction are included. The proteins are embedded in a hexagonal double membrane lipid bilayer (hydrophobic lipid tails are shown as grey sticks, lipid phosphorous atoms as orange spheres), solvated with water (cyan) and NaCl ionic bath (blue and yellow spheres).

**Figure 2.**
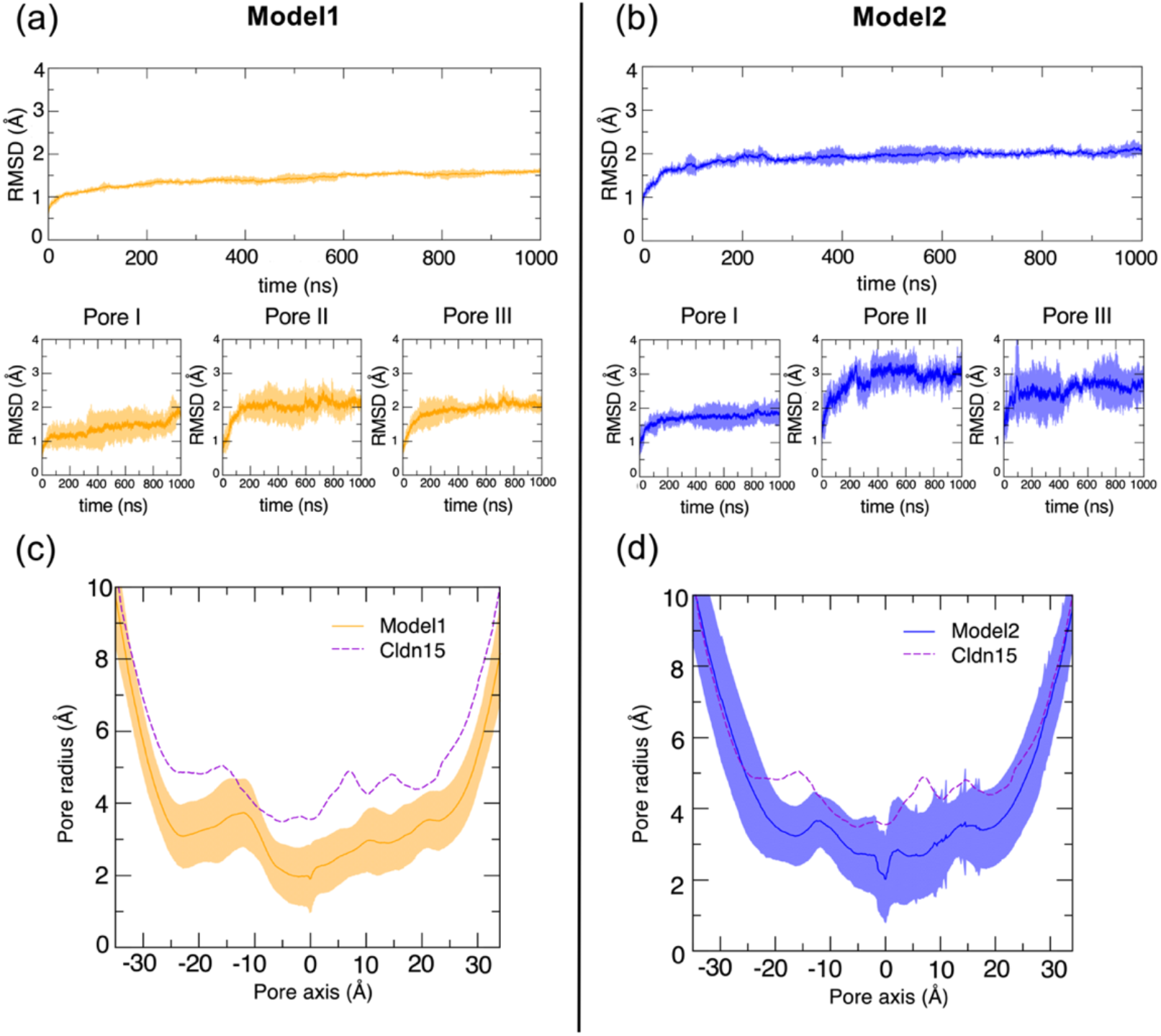
Structural analysis of the Claudin-10b triple-pore models. (**a**,**b)** Backbone RMSD of the ECL domains of Model1 and Model2, respectively. Upper panels: all paracellular domains. Lower panels: individual pores. **(c**,**d)** Pore radius profiles of Model1 and Model2, respectively, compared to that of Cldn15 multi-pore (purple dotted line). Dark lines indicate averages over each system’s replica, while shaded areas represent standard deviations.

#### Pore radius

The pore radii of paracellular channels were calculated with HOLE ^[50]^. **Figures 2c,d** show the radius profiles along the channel axis (central pore) for the Cldn10b Model1 and Model2, compared with that of Cldn15 (**Figure S1a)**. All systems are characterized by the typical hourglass shape previously observed in JDR-based architectures ^[38,43,44]^. For both Cldn10b models, the maximal pore constriction is about 2.2 Å, at the pore center, smaller than that found for the Cldn15 multi-pore architecture (**Figure 2c,d**, purple dashed line).

#### Assessment of cis-linear interactions

The simulation of multi-pore structures permits the analysis of intermolecular *cis-linear* interactions that stabilize the TJ strand ^[14,36]^. In the Cldn15 crystal structure (PDB ID: 4P79 ^[36]^), this network is mainly established by the residue M68 of one monomer fitting the hydrophobic cage arranged by residues F146, F147, and L158 of the neighboring one. These amino acids are conserved among various Cldn homologs, including Cldn10b (**Figure S1b)**. The *cis-linear* interactions were monitored by calculating inter-residue distances during standard MD simulations for both Model1 (**Figure 3a,b**) and Model2 (**Figure 3c,d**), and compared to values for the same distances from the Cldn15 crystal (**Figure 3c,d**, red dotted lines).

**Figure 3.**
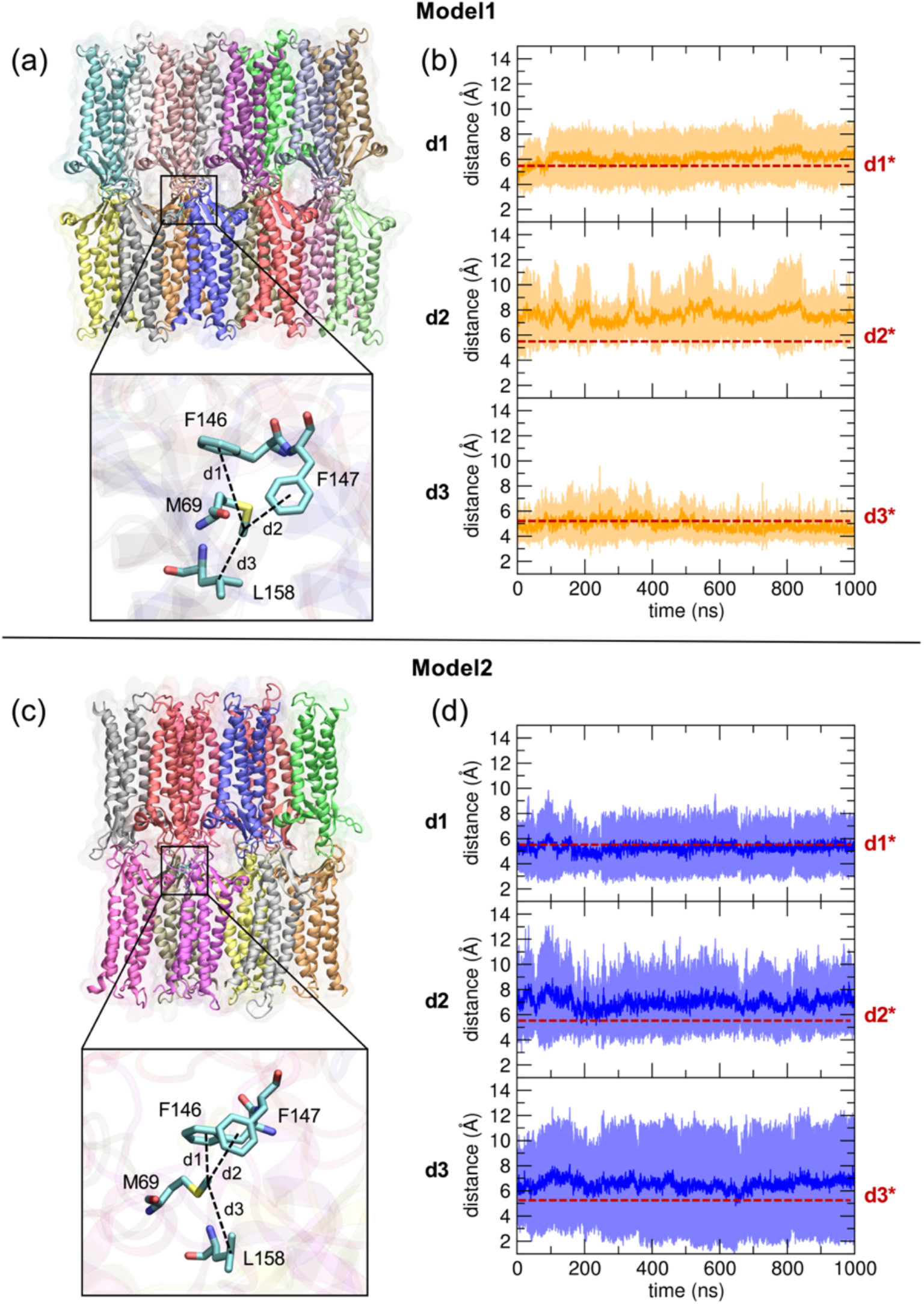
Analysis of the cis-linear interactions in the Cludin-10b triple-pore models. (**a,b)** Representation and time evolution of **the** distances between M69 and F146 (d1), F147 (d2) and L158 (d3) for Cldn10b Model1. **(c,d)** Same for Model2. Dark lines indicate averages over each system’s replica, while light areas represent standard deviations. Red dotted lines are the corresponding values in the Cldn15 crystal structure (PDB ID: 4P79).

Results are also summarized in **Table 1**. In Model1, the distances between M69 and F146 and between M69 and L158, named d1 and d3, respectively, are stable and consistent with those in the Cldn15 crystal (d1* and d3*; **Figure 3b**). The distance between M69 and F147 (d2) exhibits minimal fluctuations but differs by ∼ 2 Å from the crystal one (d2*). Similar results are found for Model2 (**Figure 3d**), where all distances are stable, d1 is consistent with d1*, but small differences (∼ 1 Å) are observed between d2 and d3 and the corresponding crystal values (d2* and d3*).

**Table 1.** Distances mapped for the Claudin-10b triple pore models. Atom names between parentheses are indicated as follows: CD is the side chain Cδ atom, CE the Cε atom, and CG the Cγ atom, while COM is the center of mass of the aromatic side chain. For distances d1, d2 and d3, corresponding to the cis-linear interactions, we report the corresponding distances from the Cldn15 crystal structure (d1*, d2* and d3* in Figure 3).

| Distance name | Residues (atoms) | analysis | Distance (Å) |  |  |
| --- | --- | --- | --- | --- | --- |
|  |  |  | Model1 | Model2 | Claudin-15 crystal (PDB ID: 4P79<sup>[36]</sup>) |
| d1 | M69 (CE) – F146 (COM) | Cis-linear interactions | 6.09 ± 0.41 | 5.26 ± 0.33 | 5.33 |
| d2 | M69 (CE) – F147 (COM) | | 7.58 ± 0.47 | 6.99 ± 0.50 | 5.29 |
| d3 | M69 (CE) – L158 (CG) | | 4.92 ± 0.40 | 6.53 ± 0.43 | 5.10 |
| d4 | D73 (CG) – D73 (CG) | Acidic pore-lining residues cross-distances | 15.71 ± 0.48 | 23.85 ± 0.96 | |
| d5 | E145 (CD) – E145 (CD) | | 21.68 ± 0.90 | 20.53 ± 0.49 | |
| d6 | E153 (CD) – E153 (CD) | | 11.17 ± 1.38 | 12.94 ± 1.27 | |
| d7 | D36 (CG) – D36 (CG) | | 28.71 ± 1.12 | 11.09 ± 0.48 | |
| d8 | D56 (CG) – D56 (CG) | | 12.86 ± 0.50 | 11.31 ± 0.78 | |

#### Analysis of the electrostatic paracellular environment

The distribution and orientation of the charged pore-lining residues govern the ion-selectivity of Cldn pores ^[51]^. Concerning Cldn10b, the paracellular cavities of both Model1 (**Figure 4a**) and Model2 (**Figure 4b**) are largely populated by acidic residues facing the lumen of each pore: halfway through each pore, in correspondence with the maximal constriction, four D56 residues, one from each monomer, form a ring and point their sidechains towards the pore; moving towards the exits of the channels, two couples of D36 and E153 are symmetrically oriented in the intermediate segments, while the D73 and E145 ones are located at the two entrances. On the other hand, the basic amino acids are located far from the pore volume accessible to ions or other permeating molecules, except for K64, found between D36 and D56, pointing towards the lumen. As a result, the electrostatic surface of both paracellular models (**Figure 4b,d**) exhibits a predominantly acidic character, with only few basic spots near the TM domains. On the contrary, no relevant hydrophobic regions are found inside these systems, so that water is unlikely to be blocked by a hydrophobic gating mechanism, contrary to what was previously suggested ^[31]^.

**Figure 4.**
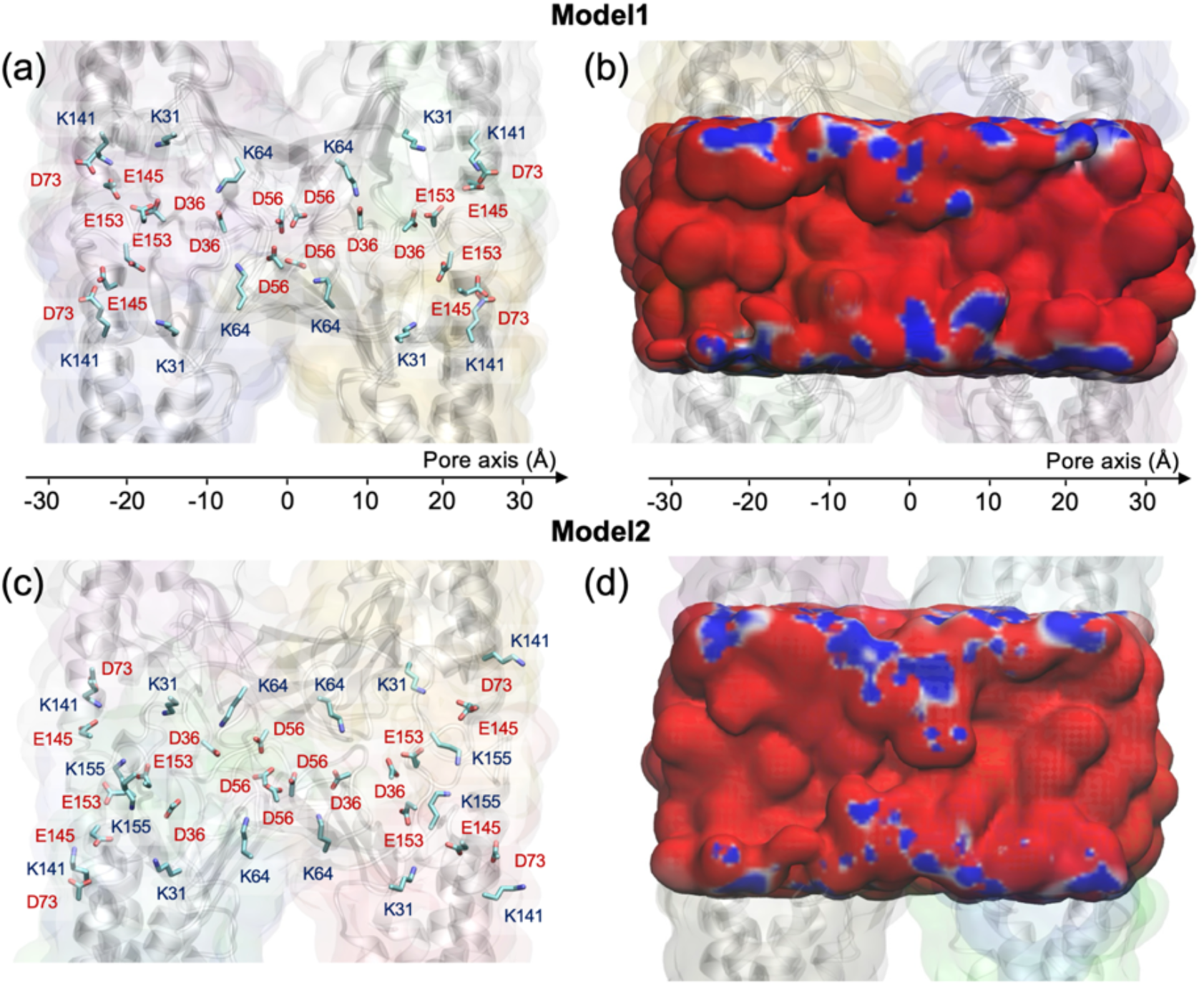
Paracellular composition of the Claudin-10b triple-pore models. (**a,b)** Orientation of charged pore-lining residues and electrostatic surface in Model1; **(c,d)** same for Model2. The electrostatic surfaces are represented with a color-code ranging from red (−5 kT/e) to white to blue (+5 kT/e).

To highlight the differences in the disposition of the acidic pore-lining residues in the two models, we calculated distances between the side chains of each pair of residues facing across the cavities (d4 to d8). Representative snapshots of Model1 and Model2 structures, highlighting the amino acids used to define the distances, are reported in **Figure 5a** and **c**, whereas **Figure 5b** and **d** show how all distances evolve over time. The major variations are found for d4 (∼8 Å) and d7 (∼17 Å), at the entrance and towards the center of the cavities, respectively, due to the dissimilar orientation of the *β*1*β*2 loops in the two Cldn10b models (**Figure S2**). Interestingly, at the constriction centers, the distances between D56 residues (d8) display only minor differences (∼1 Å). A summary of all distances discussed in this section is reported in **Table 1**.

**Figure 5.**
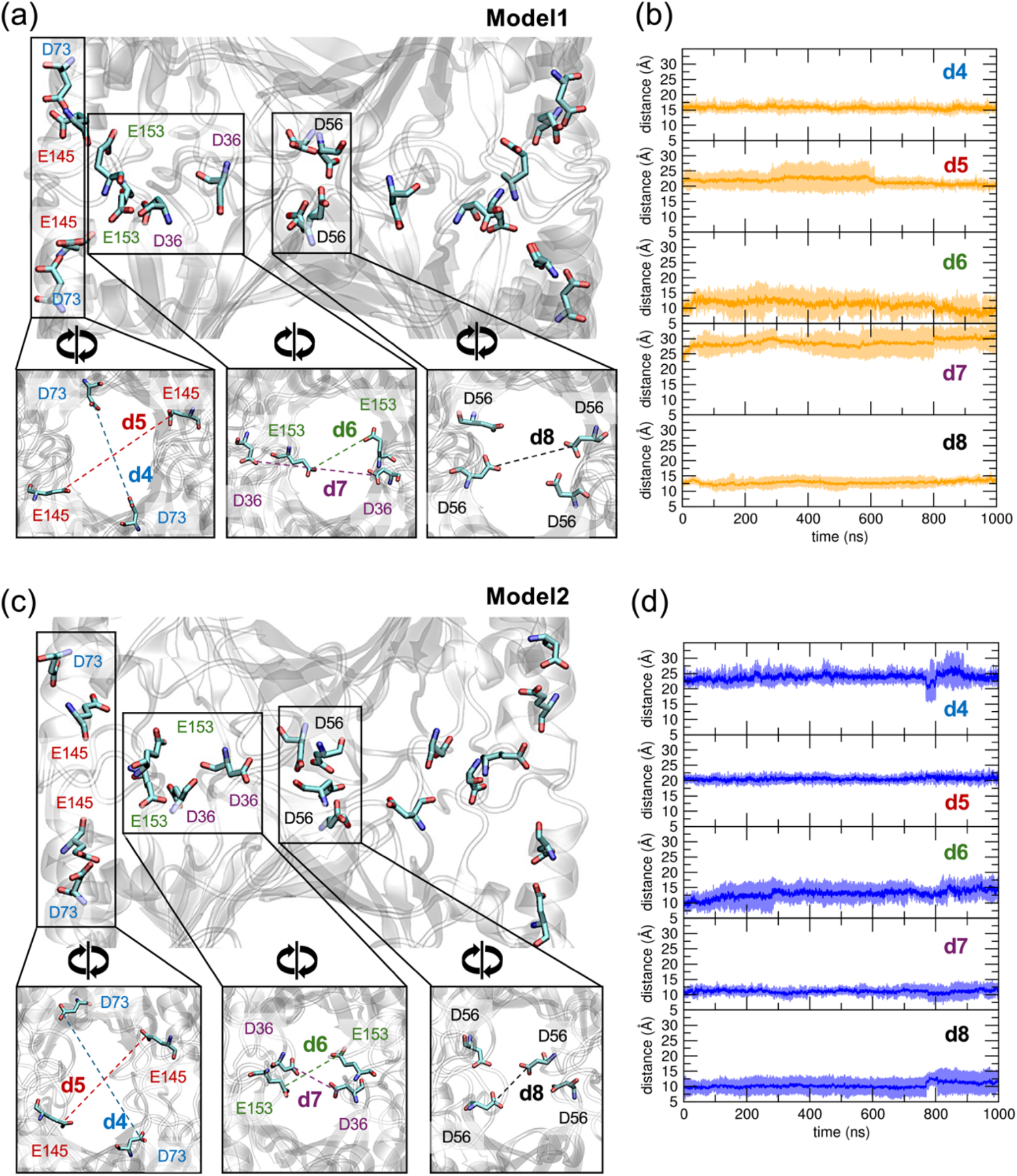
Cross-distances between opposing acidic pore-lining residues. (**a,b)** Illustration and time-evolution of distances between facing acidic amino acids in Model1: d4 (D73-D73), d5 (E145-E145), d6 (E153-E153), d7 (D36-D36) and d8 (D56-D56); **(c,d)** same for Model2. Dark lines indicate averages over all replicas and monomer pairs, while light areas represent standard deviations.

### Ion transport is mediated by pore-lining acidic sites

After the assessment of the structural properties of the two models, we analyzed the details of Na^+^ transport through the Cldn10b and Cldn15 pores. The path of Na^+^ ions visiting the paracellular central cavity of each system was monitored by recording their position along the pore axis, corresponding to the cartesian *y*-coordinate, as a function of time. Results for one replica of Cldn10b Model1 and Model2 are reported in **Figure 6a** and **b**, respectively, while other replicas are shown in **Figure S3**. In these panels, each color represents the trajectory of a single Na^+^ ion found within the pore, i.e., with a *y*-coordinate between -30 Å and 30 Å. It can be observed that most ions reside between -10 Å and +10 Å along the pore axis in Model1 and between -15 Å and +15 Å in Model2. In these channel regions, the acidic residues form coordination sites for ions to remain stably bound or interchange. Consistently, the distributions of Na^+^ ion along the pore axis, calculated using all simulated replicas (**Figure 6c**), show three peaked regions in both models, corresponding to the positions of D56 residues, in the middle, and D36 and E153 at the sides. The peaks corresponding to D36 and E153 are less pronounced in Model1 than in Model2, likely because of the more peripherical position of these residues in the former (see differences in distances d6 and d7 in **Figure 5**). To evaluate the hydration state of the permeating ions, we calculated the average number of water molecules bound to each Na^+^ ions inside the central cavity. Results indicate that the ions are coordinated by ∼ 5 water molecules on average at the pore mouths of Model1 and Model2, where they interact with D73 and E145 (**Figure S4**). On the other hand, the ions lose up to three water molecules when bound to the acidic residues within the pore (E153, D36, D56).

**Figure 6.**
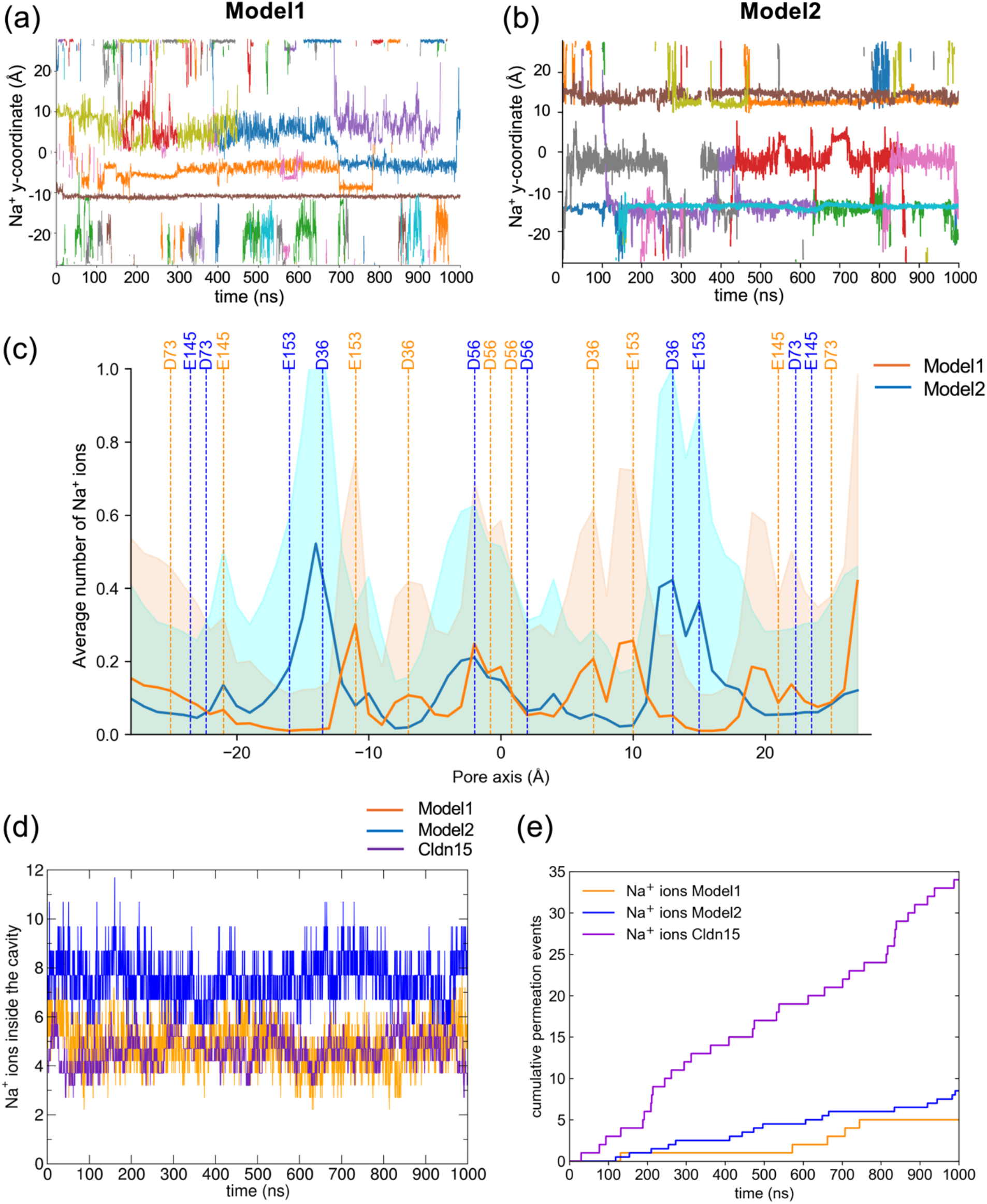
Ion conduction through multi-pore claudins architectures. (**a,b)** Time-evolution of Na^+^ positions inside the central pore of Cldn10b Model1 and Model2, respectively. Each color corresponds to a different cation. Only ions that spent at least 10% of the total time inside the pore are shown for clarity. **(c)** Distributions of Na^+^ ions along the pore axis. The profiles and the associated errors are calculated as mean and standard deviation over the three replicas of each model. Dashed lines correspond to the positions of indicated residues’ Cαatoms in the equilibrated structures. **(d)** Number of Na^+^ ions inside the pores of indicated systems along the simulated trajectories (averaged over the replicas for Cldn10b). **(e)** Cumulative number of ions that progressively crossed Cldn10b Model1 (orange), Model2 (blue), or Cldn15 (purple) pore during the MD simulations. Traversing in both directions was considered, and data for Cldn10b were averaged over replicas.

The average number of Na^+^ ions occupying the paracellular cavities during the trajectories varies in the two structures: about 5 and 8 Na^+^ ions are found inside Model1 and Model2 pores, respectively (**Figure 6d**). Since both Cldn10b and Cldn15 are known to be cation-selective, we compared these results to those for Cldn15. Despite the latter’s greater pore size, we found that the number of cations in its cavities is approximately the same as Cldn10b Model1 (∼5 Na^+^ ions). On the other hand, both Cldn10b models show a significantly lower rate of Na^+^ ions fully traversing the pores -that is, entering from one side and escaping from the other-than Cldn15 (**Figure 6e**).

### The water flux through Claudin-10b is slowed down compared to Claudin-15

We then investigated water permeation through the Cldn10b channels, and compared it with that of Cldn15. **Figure 7a** shows snapshots from simulations of the three systems, where it can be observed that the paracellular cavities are fully hydrated. However, relevant differences are found between the water fluxes through the two claudins. First, compared to Cldn15, both Cldn10b models can fit fewer water molecules in the cavity along the trajectory (**Figure 7b**). Secondly, the rate at which water molecules fully traverse the pore is substantially lower for Cldn10b than for Cldn15 (**Figure 7c**). To rule out the possibility that these water molecules are part of the cation hydration shells, we calculated the percentage of time each permeating water was coordinated to a Na^+^ ion. **Figure 7d** shows that, in the Cldn10b Model1, approximately 50% of the permeating water molecules do not interact with Na^+^ ions, while the remaining molecules bind Na^+^ for a maximum of 5% of the trajectory, and, in any case, for less than 20-30% of it. For Model2 and Cldn15, about 20% of the crossing waters are never part of the ions’ hydration shells; the remaining ones coordinate ions for less than 40% of the time and up to about 10% of the time at most.

**Figure 7.**
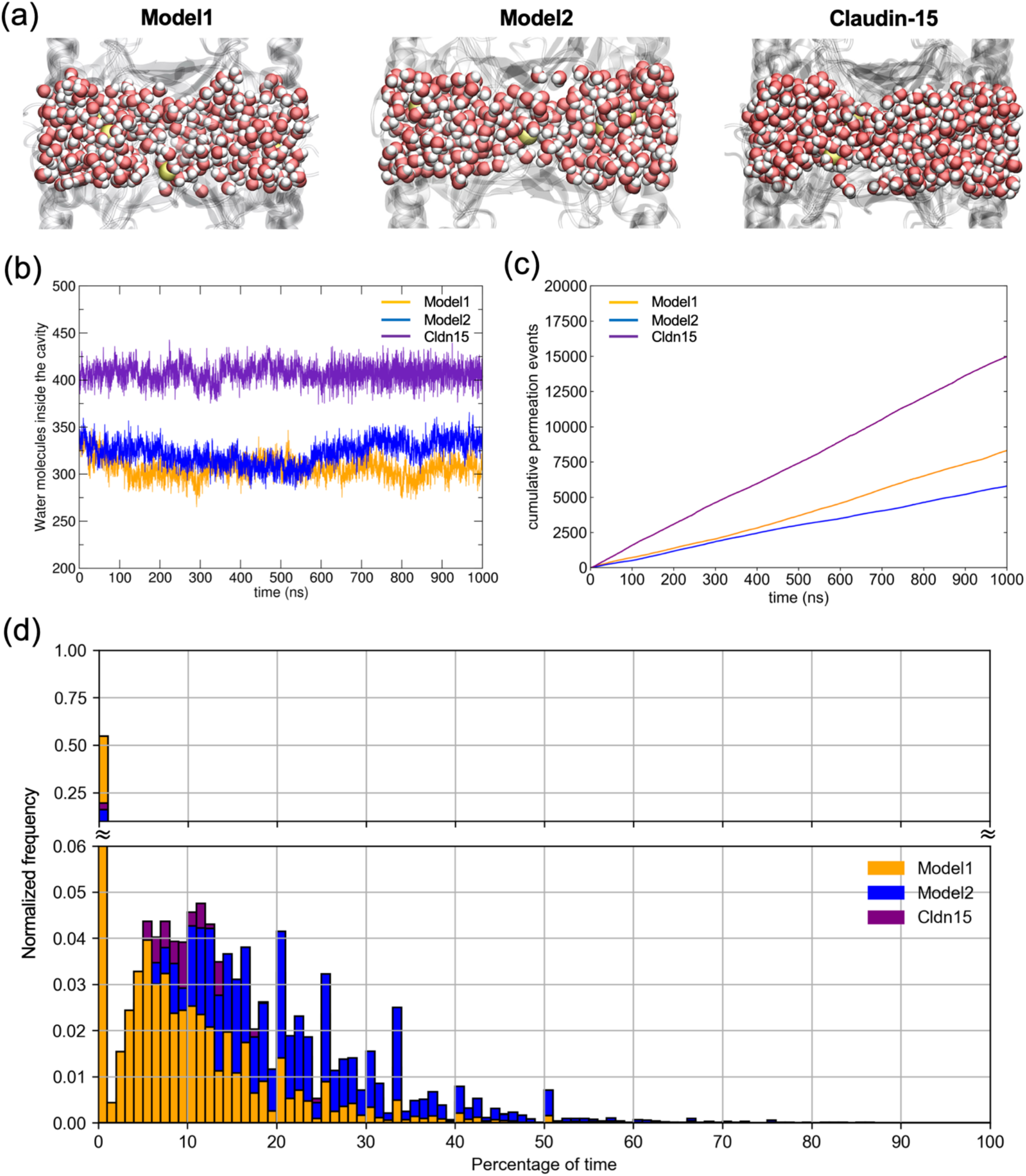
Water permeation through the Claudin-10b and Claudin-15 models. **(a)** Representative snapshots of the hydrated paracellular cavities of Cldn10b and Cldn15 models. **(b)** Average number of water molecules inside Cldn10b Model1 (orange), Model2 (blue) and Cldn15 (purple) pores. **(c)** Cumulative number of water molecules that progressively crossed the Cldn10b Model1 (orange), Model2 (blue) or Cldn15 (purple) pores during simulations. Traversing in both directions was considered, and data for Cldn10b were averaged over replicas. **(d)** Histograms of the percentage of simulated time each traversing water molecule is coordinated to a Na^+^ ion.

Overall, the standard MD simulations suggest that, although to a different extent, in the two models, Na^+^ ions populate the Cldn10b pores strongly interacting with the acidic pore-lining residues, and their permeation rate is lower than in Cldn15. The rate of translocating water molecules is also smaller than in Cldn15, and most of them traverse the pore as free diffusing molecules, being part of Na^+^ hydration shells only for small portions of the trajectories.

### Free energy calculations of ion and water permeation

To quantify the selectivity properties of Cldn10b paracellular architectures, we calculated the one-dimensional FE profiles for single ions or water molecules permeating through the pores, using the US-WHAM method ^[52,53]^. In the calculations, all ions except the tagged ones were kept outside the pore. Results show that in both Model1 (**Figure 8a**) and Model2 (**Figure 8b**) the passage of the anion is hindered by FE barriers of ∼11 kcal/mol, peaked at the central region of the pore, in correspondence with the maximal constriction and the ring of negatively charged D56 residues. Consistently, the profiles for Na^+^ and K^+^ show deep attractive minima in the same region. However, the Cl^-^ curves of the two models have different shapes, whereas those of the cations differ in shape and minimum depth. While, in Model1, the barrier to the anion is narrower, growing beyond 10 kcal/mol only between -10 Å and 10 Å, in Model2 it is much broader. The cation profiles have the same difference in width, but while the minimum reaches about -4/5 kcal/mol in Model1, it is -8 kcal/mol in Model2. These differences arise from the positions and orientation of the pairs of D36 and E153 residues in the two models: with respect to Model1, in Model2 they are positioned farther away from the middle of the pore axis, thus expanding the region of their effect; moreover, in the same model, the D36 are closer to the pore lumen and with their side chains oriented towards it (see **Figure 5**). These findings are consistent with those from standard MD simulations, which indicated that Model1 pore is less populated by Na^+^ ions than Model2 (**Figure 6d**). Finally, the FE profiles for the permeation of a single water molecule indicate a barrierless process in both models (**Figure 8a** and **b**, black line). Although not completely unexpected, given the full hydration of cavities seen in standard MD (**Figure 7a, b**), this result seems not to agree with the limited water permeation observed in the Cldn10b models (**Figure 7c**). Opposite to standard MD, however, the FE calculations were performed with no ions inside the pores, and so we set out to determine whether the profiles would change if we allowed Na^+^ ions within the cavity.

**Figure 8.**
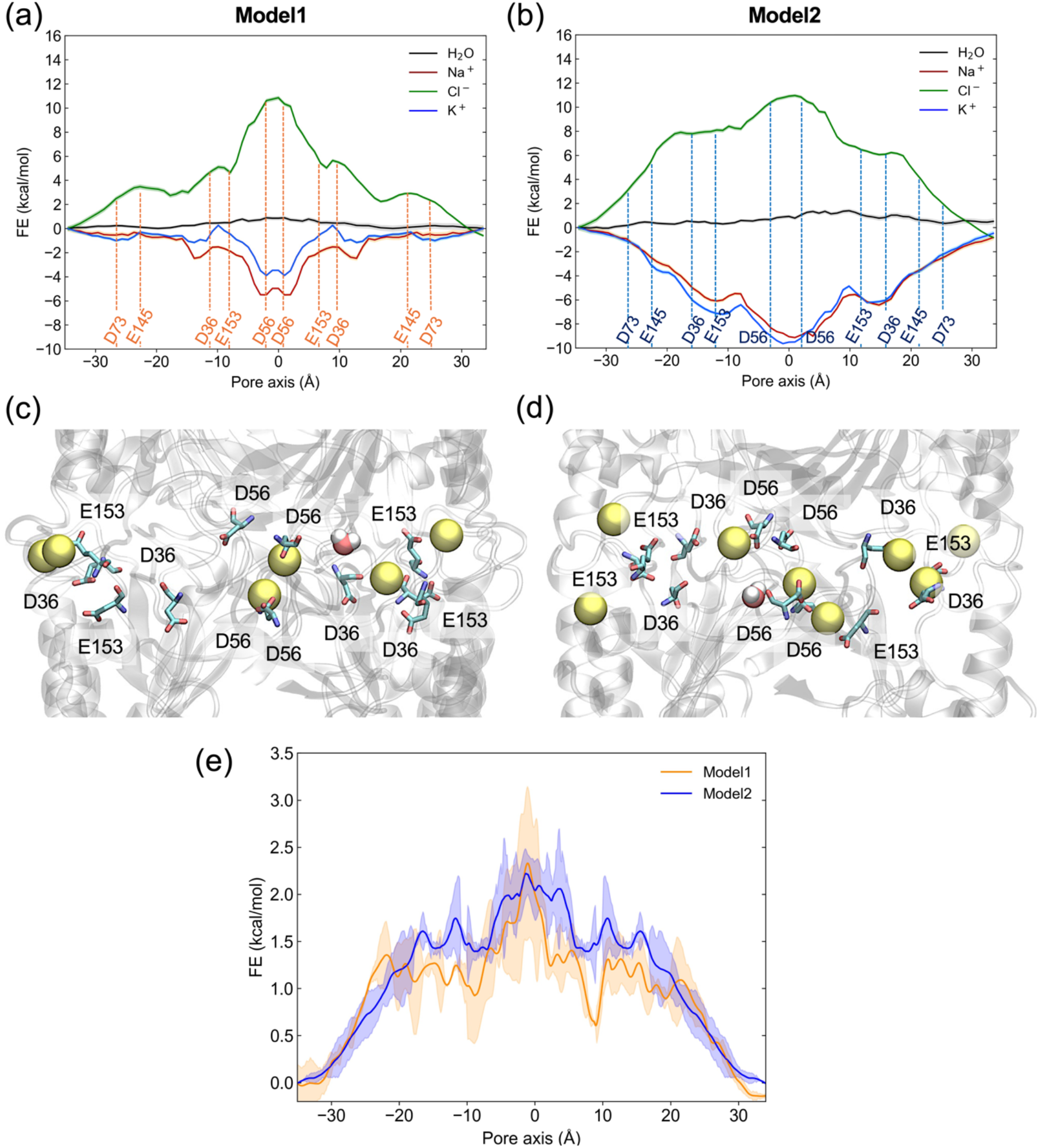
Free energy calculations of ions and water permeation through the Claudin-10b pores. **(a,b)** FE profiles for single-ion/water molecules permeation through Model1 and Model2, respectively. The positions of pore-lining acidic residues, identified based on the coordinates of their Cα atoms in the equilibrated structure, are indicated. **(c,d)** Representative snapshots of Model1 and Model2 configurations selected to compute the FE profiles of water permeation in the presence of multiple Na^+^ ions inside the pore. **(e)** FE profiles for water permeation through ion-occupied Model1 and Model2.

We then repeated the US calculations for water permeation through the pores by considering distinct realizations of bound ions, based on conformations observed in the standard MD trajectories. We started with one Na^+^ ion in Model1, at the central D56 ring (**Figure S5a**), and three Na^+^ ions in Model2, one coordinated by the central D56 residues and two by the pairs of D36 (**Figure S5b**). Results show again no barriers to water permeation, in both models with limited ion occupation (**Figure S5c**). We then increased the number of Na^+^ atoms inside the pores, using configurations representative of the average occupation of Na^+^ ions observed during the unbiased trajectories, i.e., six Na^+^ ions in Model1 (**Figure 8c**) and eight Na^+^ ions in Model2 (**Figure 8d**). The cations were confined within the pores but left free to move inside of them. These additional calculations were performed using the Temperature Accelerated Molecular Dynamics (TAMD) method ^[54]^, and resulted in symmetrical FE profiles with maxima of ∼ 2.2 kcal/mol for both Model1 and Model2, localized at the center of the pore (**Figure 8e**). These results indicate that the higher the number of Na^+^ ions inside the pore, the more limited is the passage of water, thus suggesting that the restricted permeability of Cldn10b to water is due to a blockade formed by cations within the cavity.

## Discussion

The recent endeavors to elucidate the structural organization of Cldn proteins within epithelial and endothelial TJ strands have profited from using computational methods to address both their functional ^[38–40,42–47,49,55]^ and mechanical ^[41,56,57]^ features. Most of these investigations employed the structural template introduced by Suzuki et al., ^[35]^ based on the Cldn15 crystal ^[36]^. This *joined-double row* (JDR) model has been successfully validated for many homologs with different physiological functions, including channel-forming systems such as Cldn15 ^[38,40]^, Cldn2 ^[42]^, Cldn10b ^[46]^, or Cldn4, ^[44]^ and even barrier-forming systems such as Cldn5 ^[43,55]^. In all these works, it has been shown that the electrostatic environment produced by the pore-lining residues governs the accessibility of the ions. However, little attention has been devoted to the transport of water ^[56]^. The mechanistic description of this phenomenon represents a notable challenge to these structures, since, while the size of water molecules (VdW radius ∼ 1.7 Å ^[58]^) allows passage through the pores, some Cldn subspecies are permeable to ions but not to water. Cldn10b, expressed in the thick ascending limb of the nephron’s loop of Henle, where it forms channels permeable to cations but not to water, is a remarkable example ^[31,34]^.

In this work, we used MD simulations to test whether two variants of the JDR structural model for paracellular TJs can recapitulate the selectivity properties of Cldn10b. Both variants comprise three pores and represent a realistic proxy of the multimeric organization of Cldn proteins in TJs, including interactions stabilizing the aggregates within the same cell and across the paracellular cleft. However, they differ in the orientation of the segment connecting the *β*1 and *β*2 strands of each monomer’s ECL1 (the *β*1*β*2 loop), which was missing from the original JDR architecture and had to be modeled. Since different loop conformations result in distinct orientations of residues that are known to be relevant for Cldn selectivity, it was questioned whether both models could reproduce the functional properties of paracellular channels.

Our MD simulations on the microsecond timescale indicate that the two conformations are stable and have a comparable pore shape and size, with a minimal radius of about 2.2 Å at the center. This is smaller than those found in all multi-pore systems modeled so far for channel-forming Cldns, such as Cldn2 ^[42,48,59]^ (minimal radius ∼ 3.4 Å), Cldn4 ^[44,48]^ (∼ 3.2 Å), and Cldn15 ^[39,48,60]^ (∼3.8 Å), suggesting that the steric factor plays a critical role in the regulation of paracellular transport. In particular, the difference with Cldn15 is in line with previous computational ^[35,39,40,47]^ and experimental data ^[19,32,33,45]^.

Both models show multiple Na^+^ binding sites within the pores (corresponding to residues D56, D36, and E153), among which ions can exchange while shedding part of their hydration shell. The D56 ring, previously identified as the selectivity filter ^[34,45]^, is located at the central and narrowest region of the pore. The only basic residue lining the pore axis is K64 that, as previously suggested ^[47]^, may compete with cations for interactions with acidic residues, establishing transient contacts with the neighboring D36 or D56. The cation-selective property of Cldn10b was reproduced for both architectures using FE calculations of single-ion permeation. Although the actual transport mechanism results from the crossing of multiple ions, the single-ion FE captures the fundamental features of ion-protein interaction as, for example, the location of binding sites and repulsive barriers. Our data show that both models generate FE barriers to the passage of anions while being attractive for cations. However, while the distinct *β*1*β*2 loop conformations do not affect the height of the barriers, they result in different profile shapes and minima depths: in the structure from Piontek and collaborators (Model2), the curves are broader, and the minima are almost twice as deep as in Model1. This is due to the D36 residues being closer to the pore axis, with their side chains pointing towards the lumen. The higher attractivity of Model2 for cations results in about 8 Na^+^ ions found on average inside the pore, whereas in Model1 and Cldn15 they are around 5. This notwithstanding, the flux of cations across both Cldn10b structures is much smaller than in Cldn15. This might be related to the largest pore size of Cldn15, which avoids jamming of ions. Finally, water permeation analysis reveals that Cldn10b channels can accommodate approximately 20% fewer water molecules than the wider Cldn15 ones, with a flux across the pore that is 2-3 times smaller. The mechanism of water crossing, however, is different between the two models, with more free molecules traversing the pore in Model1 than in Model2. This might be a consequence of the smaller number of ions inside Model1 pore, resulting in fewer interactions with water.

Altogether, our data demonstrate that both Cldn10b pore models are cation-selective, albeit to a different extent, and that they are less permeable to water compared to Cldn15, consistent with the experimental evidence ^[31,32,34]^. This behavior might result from the combination of the strongly acidic character and the narrow size of the Cldn10b pores, causing jamming of cations that enter the cavity and oppose water passage. This mechanism confirms previous hypotheses, ^[47]^ and is supported by our FE calculations, which showed increased FE barriers for water as Na^+^ ions crowded the pore. The differential selectivity for ions and water represents a challenge for computational and experimental investigations. To date, theoretical studies have mostly been devoted to canonical transmembrane channels, in which the protein is embedded in the hydrophobic lipid environment ^[61–70]^. Some works ^[71–74]^ showed that, in narrow pores, apolar residues might create dry regions that could stop the flow of water and ions even without mechanical closure of the pathway (hydrophobic gating). A similar mechanism was originally supposed for Cldn10b,^[31]^ but the hydrophilic environment of the pore and the flat FE profiles obtained for single-water molecule calculations do not support this hypothesis.

In summary, our data show that the JDR model, even with different conformations of the ECLs, is compatible with Cldn channels that are selective for ions and not for water, further expanding its validity as representative of TJ assemblies. This study offers insights into the molecular mechanism underpinning paracellular water selectivity, a crucial component of epithelial transport systems.

However, the small FE barriers and the large number of solvent molecules found in the paracellular space do not fully agree with null water transport. This may be due to the partial description offered by the atomistic TJ models, since they do not encompass multiple strands, as observed in tight epithelia ^[75]^, or long time-scale conformational transitions eventually closing the pores. Moreover, no concentration gradient is present between the two sides of our paracellular systems. Future investigations would benefit from the simulation of larger oligomers and for longer times, outlining a more detailed picture of the junctional paracellular architecture.

## Materials and Methods

### Construction of the Claudin-10b and Claudin-15 multi-pore models

We built two distinct Cldn10b JDR models, each comprising three adjacent pores. The first one, Model1, was generated using as a template an equilibrated configuration of the Cldn15 double-pore system we produced before ^[39]^. From this model, we extracted four Cldn15 monomers forming the interface between two adjacent pores (**Figure S6**). One of the Cldn15 units was used as a template to model the Cldn10b monomer with SWISS-MODEL ^[76]^. Then, four replicas of the resulting monomer were superimposed on the Cldn15 ones. Major clashes were solved using GalaxyRefineComplex ^[77]^, and the resulting configuration was replicated to form a dodecameric triple-pore system with the help of VMD 1.9.4 ^[78]^. Finally, the four most external protomers were added to preserve the *cis*-interactions of each protomer. The final hexadecameric multimer was relaxed with the Generalized-Born Implicit Solvent (GBIS) method ^[79,80]^. The cutoff for the VdW interactions was set to 14 Å and the time step was 2 fs. After a starting minimization, 15 ns of equilibration were performed with NAMD3 ^[81]^ and the CHARMM36m ^[82]^ force field.

The second Cldn10b triple-pore model, here named Model2, corresponds to the *octameric interlocked barrel* configuration resulting after 100 ns of MD simulations produced by Prof. J. Piontek and collaborators, published in Ref. ^[47]^ The Cldn15 multi-pore system was generated via the same procedure of Model1.

### Assembly of the Systems

Each Cldn10b and Cldn15 multi-pore structure was translated so that its center of mass coincided with the origin of the reference system in VMD1.9.4 ^[78]^, and oriented with the pore axis parallel to the cartesian *y*-axis. Two hexagonal membranes of pure 1-palmitoyl-2-oleoyl-glycero-3-phosphocholine (POPC) were generated using the *membrane builder* tool of CHARMM-GUI ^[83,84]^ and equilibrated separately for 10 ns with the NAMD3 software ^[81]^ and the CHARMM36 ^[82]^ force field using hexagonal Periodic Boundary Conditions (PBC). Then, the complexes were embedded in the equilibrated double membrane bilayer and solvated with TIP3P water molecules ^[85]^ and a physiological NaCl ionic bath (0.15 M). The final simulation boxes are hexagonal prisms with a base inscribed in a square of approximately 200.0 × 200.0 Å^2^ and a height of about 160.0 Å, counting ∼ 500,000 atoms (**Figure 1**). The topology file was built with the *psfgen* tool of VMD 1.9.4 ^[78]^, using the parameters of the CHARMM36 ^[82]^/ CHARMM36m ^[86–88]^ force field. Disulfide bridges between residues C53 and C63 fin the ECL1 of each protomer were preserved for both Cldn10b and Cldn15 proteins.

### Equilibration and standard MD simulations

All systems were equilibrated with a multi-step protocol comprising a progressive release of harmonic restraints on heavy atoms (**Table S1**). After first energy minimization, 100 ns of equilibration and 1000 ns of production were carried out in the NPT ensemble at a constant temperature and pressure of 310 K and 1 bar, respectively, maintained by a Langevin thermostat and Nosé-Hoover Langevin piston ^[89,90]^. Following the setup suggested by the CHARMM-GUI configuration files for NAMD ^[83]^, the oscillation piston period was set to 50.0 fs and the damping time scale to 25.0 fs. The damping coefficient of the Langevin thermostat was set to 1 ps^-1^. Electrostatic and van der Waals (VdW) interactions were calculated with a cutoff of 12 Å as customary with the CHARMM force field. A switching function was applied starting to take effect at 10 Å to obtain a smooth decay ^[91]^. Long-range electrostatic interactions were computed using the Particle-Mesh Ewald (PME) algorithm ^[92]^, with spline interpolation order 6. A maximum space between grid points of 1.0 Å was used. Chemical bonds between hydrogen atoms and heavy atoms were constrained with SHAKE ^[93]^, while those of the water molecules were kept fixed with SETTLE ^[94]^. A time step of 1 fs was employed for the first 20 ns of equilibration. Then, it was increased to 2 fs for the subsequent 80 ns. A summary of the restraints and time steps adopted during each phase of the equilibration is reported in **Table S1**. The production phase was conducted maintaining the same restraints on the Cαatoms belonging to the TM helices as the last equilibration step, mimicking the constraining effect of additional, adjacent protomers in the physiological strands. One replica of each Cldn10b system and the one for Cldn15 were simulated using a timestep of 2 fs. Two other replicas of each Cldn10b model were simulated using the hydrogen-mass repartitioning (HMR) method ^[95–97]^, adopting a time step of 4 fs. In HMR, the mass of hydrogen atoms not belonging to water molecules is adjusted to 3.024 amu, and those of the covalently bound heavy atoms are proportionally scaled down so that the total mass of the system is conserved. In these simulations, as prescribed in Ref. ^[95]^, the oscillation period of the Langevin piston period was set to 300 fs, while the damping time scale was 150 fs. MD simulations were performed with NAMD3 ^[81]^ and the CHARMM36m ^[86–88]^ parameters for the lipid and the protein, respectively, together with the TIP3P model for water molecules ^[85]^ and the associated ionic parameters with NBFIX corrections ^[98–100]^.

### Analysis of the standard MD simulations

#### Root-mean square deviation

We calculated the root-mean square deviation (RMSD) of the backbone atoms of all paracellular domains formed by the Cldn10b ECLs (residues 28 to 73 and 149 to 161 of each monomer). The analysis was conducted along the simulated trajectories of each replica with the help of VMD 1.9.4 ^[78]^ and the Tcl scripting interface. For each Cldn10b model, RMSD average values and errors were calculated using the three replicated simulations.

#### Pore radius

The size of the paracellular pores was calculated using HOLE ^[50,101]^. This program maps the radius of a protein cavity along a given axis (the *y*-axis, in this case) by fitting a spherical probe in the space not occupied by the VdW spheres of the pore-lining atoms. A max value of 15 Å was chosen as a threshold for all systems. Radius values of each Cldn10b system were calculated for the central pore every 50 ns of the replicated trajectories.

#### Electrostatic surface

Electrostatic potential maps were computed with the adaptive Poisson-Boltzmann solver (APBS) code ^[102]^, using the default parameters set by the developers.

*Cross-distances*. Cross-distances between representative residues of each system were calculated with the Tcl scripting interface of VMD 1.9.4 ^[78]^. The distances are calculated by considering the most external C-atom of the sidechain in the case of aliphatic amino acids, or the center of mass of the benzene ring in the case of aromatic residues.

#### Number of water/ion permeation events

The number of water molecules or ions crossing the central paracellular cavity during the simulated trajectories was calculated with VMD 1.9.4 ^[78]^ and an *in-house* Tcl script that tags each particle entering the pore, registers its *y*-coordinate and counts a permeation event when it crosses the cavity exiting from the opposite side. This means that a permeation event was counted when an ion or water molecule with a *y*-coordinate < -30 Å at a given timestep t1 passed through the central pore and reached a new position with a *y*-coordinate > 30 Å at a second instant t2 > t1. The calculation was repeated to consider permeation events occurring in both directions through the central pore.

#### Ion hydration sphere

The number of water molecules in the Na^+^ hydration shell was calculated with VMD 1.9.4 ^[78]^ and the Tcl scripting interface by setting a cutoff to the distance between the ion and the water’s oxygen atom of 3.0 Å, in agreement with the value determined in Ref. ^[103]^.

### Free Energy calculations

#### Umbrella Sampling simulations

The free energy (FE) profiles of single water molecules and single ion permeation were calculated with the Umbrella Sampling (US) ^[52]^ method, with the same setup we adopted in previous works ^[38,39,43,44,55]^ and the Colvars module ^[104]^. Accordingly, one single collective variable (CV) is considered, represented by the *y*-coordinate of the tagged ion or water molecule that permeates the central paracellular cavity. A summary of the US simulations is reported in **Section S1**. Before the production phase, each US window was energy-minimized and equilibrated in two steps, performing 1 ns with a time step of 2 fs, followed by 1 ns with a time step of 4 fs and the HMR setup. Then, each window was simulated under HMR, with all details specified for the standard MD simulations, until convergence was achieved. For water, 40 ns per window was required, while 60 ns and 80 ns per window were carried out for both cations and Cl^-^, respectively, resulting in a total of ∼ 17 μs of accrued simulation time per each Cldn10b model. In the case of the calculations of water permeation with one (for Model1) or three (for Model2) Na^+^ ions coordinated inside the cavity, each cation was restrained between opposing pairs of acidic residues (one D56 pair for Model1, two D36 pairs and one D56 pair for Model2) as shown in **Figure S5A** and **S5B**. We used the Colvars module to implement the half-harmonic restraints with constants of 1 kcal/(mol. Å^2^) starting to take effect at a distance between the ion and both the two facing carboxylic sidechains > 8 Å. For these calculations, each window was simulated for 60 ns, corresponding to a total of ∼ 4.2 μs of accrued simulation time per each Cldn10b model. Finally, the full FE profiles were obtained by combining the CV distributions of all windows using the weighted-histogram analysis method (WHAM) ^[105–107]^. To this aim, we employed the code from the Grossfield group available at http://membrane.urmc.rochester.edu/content/wham.

#### Temperature-accelerated Molecular Dynamics simulations

The FE profiles of water permeation through the cavities of Cldn10b triple-pores occupied by the number of Na^+^ ions suggested by standard MD simulations were computed using temperature-accelerated molecular dynamics (TAMD) ^[54,108]^. A detailed description of TAMD is provided in **Section S3**. As in the case of US calculations, the chosen CV is the *y*-coordinate of the tagged water molecule passing through the central cavity of the system. The CV-axis was split into three windows spanning from -35 Å to -10 Å, from -10 Å to 10 Å, and from 10 Å to 34 Å. Each window was equilibrated with the same procedure used for the US calculations and simulated for 200 ns under the same conditions as the standard MD simulations and adopting the HMR setup. An effective temperature 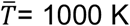 and an effective friction 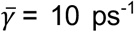 were used for to the CV’s dynamics. The FE calculations were performed by constraining six or eight ions, according to the model, inside the central cavity of the Cldn10b triple-pores with the Colvars module ^[104]^. We adopted a harmonic constant of 10 kcal/mol starting to take effect at the extremities of the pore, for *y* < -35 Å and *y* > 34 Å. The instantaneous force on the CV was collected in bins of 0.1 Å, and the FE was computed by integrating the average force with the corrected z-averaged (CZAR) estimator ^[109]^. The resulting profile and the associated error were reported as the mean and the standard deviation from two replicas. To assess the consistency between the results produced by US-WHAM and TAMD calculations, we repeated the single-ion calculations of Na^+^ permeation through the Model2 central cavity by adopting the setup described above. The comparison between the FE curves obtained with the two methods is reported in **Figure S7**.

## CRediT authorship contribution statement

**Alessandro Berselli**: Data curation, Formal analysis, Conceptualization, Investigation, Visualization, Writing – original draft, Writing – review & editing. **Giulio Alberini**: Formal analysis, Conceptualization, Investigation, Supervision, Project administration, Writing – review & editing. **Fabio Benfenati**: Supervision, Writing – review & editing, Funding acquisition and resources. **Luca Maragliano**: Formal analysis, Conceptualization, Funding acquisition and resources, Investigation, Project administration, Supervision, Writing – review & editing.

## Acknowledgments

We are grateful to Prof. Jörg Piontek and Dr. Santhosh Kumar Nagarajan for providing us their Cldn10b triple-pore model, here named Model2, and for many fruitful discussions. We acknowledge the HPC infrastructure and the Support Team at Fondazione Istituto Italiano di Tecnologia. In particular, we thank Alessia Vignolo, Sergio Decherchi and Mattia Pini for their kind assistance. We also thank Diego Moruzzo, Ilaria Dallorto, Rossana Ciancio and Arta Mehilli for administrative assistance and technical help. We acknowledge CINECA awards under the ISCRA initiative, for the availability of high-performance computing resources and support.

## Ethic statements

In this work, no animal or human study is presented.

## List of abbreviations

APBS: Adaptive Poisson-Boltzmann solver
Cldn: Claudin
COM: Center of mass
CV: Collective variable
CZAR: Corrected z-averaged
ECH: Extracellular helix
ECL: Extracellular loop
FE: Free energy
GBIS: Generalized-Born implicit solvent
HMR: Hydrogen-mass repartitioning
JDR: Joined-double row
MD: Molecular dynamics
PBC: Periodic boundary conditions
PME: Particle-mesh Ewald
POPC: 1-palmitoyl-2-oleoyl-sn-glycero-3-phosphocholine
RMSD: Root-mean square deviation
TAL: Thick ascending limb
TAMD: Temperature-accelerated molecular dynamics
TJ: Tight junctions
TM: Transmembrane helix
US: Umbrella sampling
VdW: Van der Waals
WHAM: Weighted-histogram analysis method

