## Supplementary material for "Ion and water permeation through Claudin-10b paracellular channels": SI

### Supplementary Methods

#### S1. Additional details for US simulations

Let us indicate with $\boldsymbol{\theta}(\boldsymbol{x})$ a set of CVs, i.e. functions of the atomic coordinates $\boldsymbol{x}$. In our FE calculations, we use as single CV the position of the ion or water molecule along the pore axis, oriented parallel to the *y*-axis, i.e. $\boldsymbol{\theta}(\boldsymbol{x}) = y$ . In US, the full range of the CV is split into a number of intervals, named *windows.* In each window $i$, an independent simulation is performed by adding to the MD force-field a harmonic restraining potential $U_i(y)$, confining the CV around a reference value ($y_i^0$), called *center*:

$$U_i(y) = \frac{1}{2}k(y - y_i^0)^2 \tag{S1}$$

The harmonic constant $k$ is chosen to ensure overlap of the CV distributions from adjacent windows. In our case, $k$ = 2.0 $kcal/(mol \cdot Å^2)$ for all simulations, and the pore axis was split into 70 windows, equally spaced by 1 Å. The conformation resulting after 250 ns of standard MD simulation was used as starting structure in all US windows, and the CV ion/water molecule was manually positioned at each center $y_i^0$ with the help of VMD 1.9.4 [1]. Moreover, in each window, the displacement of the ion in the direction orthogonal to the pore axis is confined within a radius $r_0 + \delta$ , where $r_0$ is the pore radius as determined by the HOLE program [2,3] and $\delta$= 2 Å.

#### S2. Methodological details for TAMD simulations

In Temperature-Accelerated Molecular Dynamics (TAMD), the sampling of infrequent events is achieved by accelerating the dynamics of a set of *M* auxiliary variables, $\boldsymbol{z}$, harmonically restrained to *M* CVs. That is, the extended system $(\boldsymbol{x}, \boldsymbol{z})$ is introduced, subjected to the following potential:

$$U_k(\boldsymbol{x}, \boldsymbol{z}) = V(\boldsymbol{x}) + \frac{1}{2}k\sum_{\alpha}^{M}(\theta_\alpha(\boldsymbol{x}) - z_\alpha)^2 \tag{S2}$$

where $V(\boldsymbol{x})$ is the MD force-field and $k > 0$ is a constant. As a consequence, the evolution of the extended system can be described by a set of equations such as for example:

$$\begin{cases} m\ddot{\boldsymbol{x}} = -\nabla V(\boldsymbol{x}) - k\sum_{\alpha}^{M}(\theta_\alpha(\boldsymbol{x}) - z_\alpha)\nabla\theta_\alpha(\boldsymbol{x}) + thermostat\ at\ temperature\ T & \text{(S3)} \\ \bar{\gamma}\dot{\boldsymbol{z}} = k(\theta(\boldsymbol{x}) - \boldsymbol{z}) + thermostat\ at\ temperature\ \bar{T} & \text{(S4)} \end{cases}$$

where $m$ is the mass matrix, $\bar{\gamma}$ is an artificial friction coefficient and $\bar{T}$ is an artificial temperature. The main idea of TAMD is that by adjusting $\bar{\gamma}$ so that the $\boldsymbol{z}(t)$ evolve slower than the $\boldsymbol{x}(t)$, and by tuning the parameter $k$ so that $\boldsymbol{z}(t) \sim \boldsymbol{\theta}(\boldsymbol{x}(t))$, one can obtain a trajectory $\boldsymbol{z}(t)$ that moves at the artificial temperature $\bar{T}$ on the free energy surface calculated at the physical $T$ [4,5]. Indeed, under such assumptions, the force that drives each auxiliary variable $z_\alpha$ in eq. (S4) satisfies:

$$k(\theta_\alpha(\boldsymbol{x}) - z_\alpha) \approx -\frac{\partial G(\boldsymbol{z})}{\partial z_\alpha} \tag{S5}$$

where $G(\boldsymbol{z})$ is the FE defined at the physical temperature $T$. Therefore, by choosing $\bar{T} > T$, the system will rapidly visit regions where FE is relatively low, and overcome barriers that would take a long time to be crossed at the physical $T$.

(a)

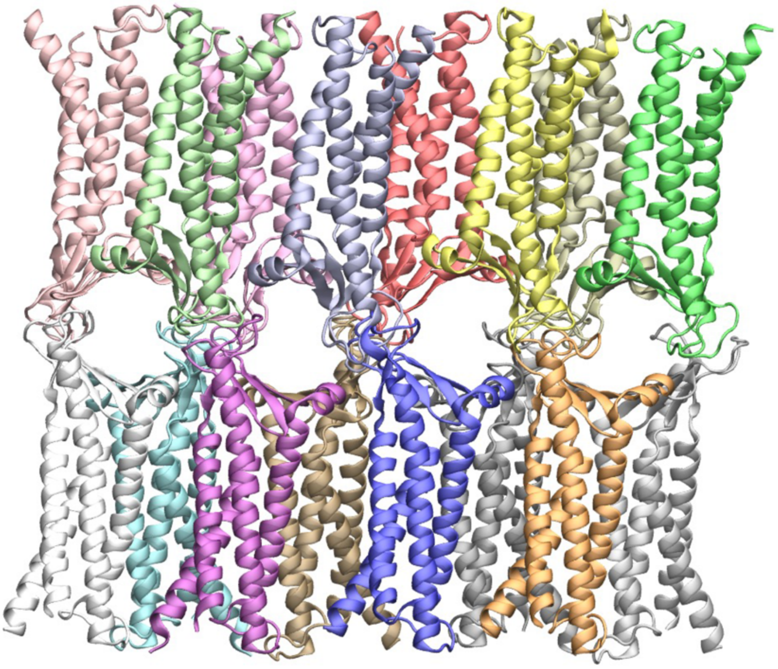

**Claudin-15**

(b)

```
CLUSTAL O(1.2.4) multiple sequence alignment


sp|P78369|CLD10b_HUMAN      MASTASEIIAFMVSISGWVLVSSTLPTDYWKVSTI-DGTVITTATYWANLWKACVTDSTG        59
sp|Q9Z0S5|CLD15_MOUSE       -MSVAVETFGFFMSALGLLMLGLTLSNSYWRVSTV-HGNVITTNTIFENLWYSCATDSLG        58
sp|P56746|CLD15_HUMAN       -MSMAVETFGFFMATVGLLMLGVTLPNSYWRVSTV-HGNVITTNTIFENLWFSCATDSLG        58
sp|P57739|CLD2_HUMAN        MASLGLQLVGYILGLLGLLGTLVAMLLPSWKTSSYVGASIVTAVGFSKGLWMECATHSTG        60
sp|O00501|CLD5_HUMAN        MGSAALEILGLVLCLVGWGGLILACGLPMWQVTAFLDHNIVTAQTTWKGLWMSCVVQSTG        60
sp|O14493|CLD4_HUMAN        MASMGLQVMGIALAVLGWLAVMLCCALPMWRVTAFIGSNIVTSQTIWEGLWMNCVVQSTG        60
                              * . : ..  :   *            *:.::    .::*:     .**  *...* *

sp|P78369|CLD10b_HUMAN      VSNCKDFPSMLALDGYIQACRGLMIAAVSLGFFGSIFALFGMKCTKVGGSDKA-KAKIAC        118
sp|Q9Z0S5|CLD15_MOUSE       VSNCWDFPSMLALSGYVQGCRALMITAILLGFLGLFLGMVGLRCTNVGNMDLSKKAKLLA        118
sp|P56746|CLD15_HUMAN       VYNCWEFPSMLALSGYIQACRALMITAILLGFLGLLLGIAGLRCTNIGGLELSRKAKLAA        118
sp|P57739|CLD2_HUMAN        ITQCDIYSTLLGLPADIQAAQAMMVTSSAISSLACIISVVGMRCTVFCQESRA-KDRVAV        119
sp|O00501|CLD5_HUMAN        HMQCKVYDSVLALSTEVQAARALTVSAVLLAFVALFVTLAGAQCTTCVAPGPA-KARVAL        119
sp|O14493|CLD4_HUMAN        QMQCKVYDSLLALPQDLQAARALVIISIIVAALGVLLSVVGGKCTNCLEDESA-KAKTMI        119
                              :*  : ::*.*   :*..:.: : :  :. .. :. : * :**       : * :

sp|P78369|CLD10b_HUMAN      LAGIVFILSGLCSMTGCSLYANKITTEFFDPLFV-EQKYELGAALFIGWAGASLCIIGGV        177
sp|Q9Z0S5|CLD15_MOUSE       IAGTLHILAGACGMVAISWYAVNITTDFFNPLYA-GTKYELGPALYLGWSASLLSILGGI        177
sp|P56746|CLD15_HUMAN       TAGALHILAGICGMVAISWYAFNITRDFFDPLYP-GTKYELGPALYLGWSASLISILGGL        177
sp|P57739|CLD2_HUMAN        AGGVFFILGGLLGFIPVAWNLHGILRDFYSPLVPDSMKFEIGEALYLGIISSLFSLIAGI        179
sp|O00501|CLD5_HUMAN        TGGVLYLFCGLLALVPLCWFANIVVREFYDPSVPVSQKYELGAALYIGWAATALLMVGGC        179
sp|O14493|CLD4_HUMAN        VAGVVFLLAGLMVIVPVSWTAHNIIQDFYNPLVASGQKREMGASLYVGWAASGLLLLGGG        179
                             .*  ..:: *   :   .     :  :*:.*      * *:* :*::*  .: : ::.*
```

***Figure S1. Claudin-15 tetrapore model and multiple sequence alignment of different Claudin homologs****.* *(a) Cldn15 multi-pore configuration based on the JDR model and composed of sixteen monomers, distinguished by color. (b) Multiple sequence alignment among human Cldn10b, mouse and Cldn15, human Cldn2, Cldn5 and Cldn4 performed with Clustal (http://www.clustal.org). Conserved amino acids stabilizing the cis-linear interactions (M69, F146, F147, L158, according to the Cldn10b numbering) are highlighted.*

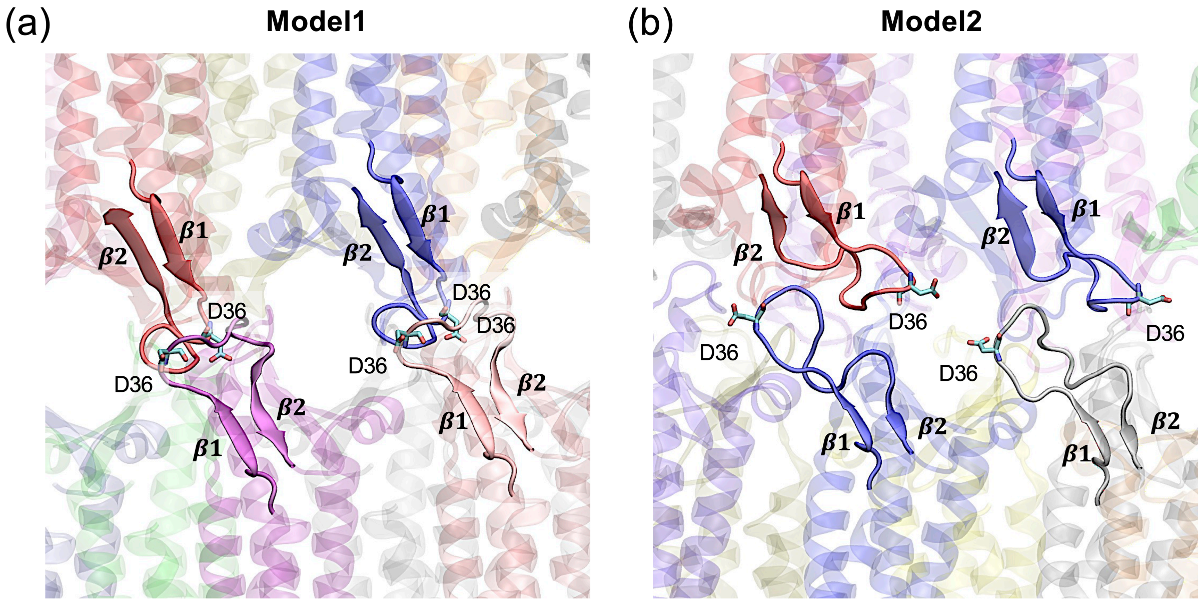


***Figure S2. Orientation of the β1β2 loop motif in the two Claudin-10b triple-pore models.*** *The different arrangement of the ECL1 β1 β2 loop in Model1 (a) and Model2 (b) is highlighted for the central pore of each system. Different Cldn10b monomers are distinguished by color, and the orientation of the D36 residues at the tip of the loop is shown.*

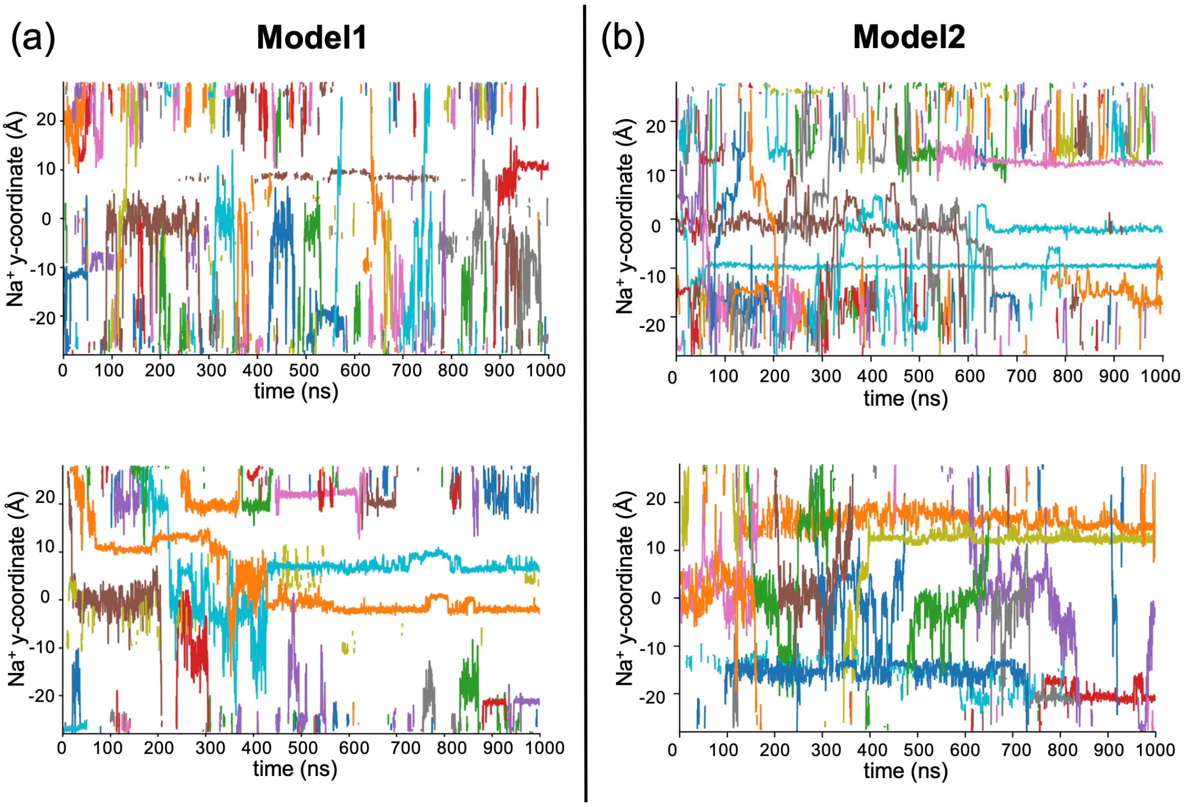


***Figure S3. Ion pathways inside the Claudin-10b paracellular cavity.*** *Each trace represents the y-coordinate, corresponding to the position along the pore-axis, of $Na^+$ ions passing through the central cavity of Model1 (a) and Model2 (b). Only ions that spent at least 10% of the total time inside the pore are shown for clarity. Different ions are distinguished by color. Each panel is representative of a replica.*

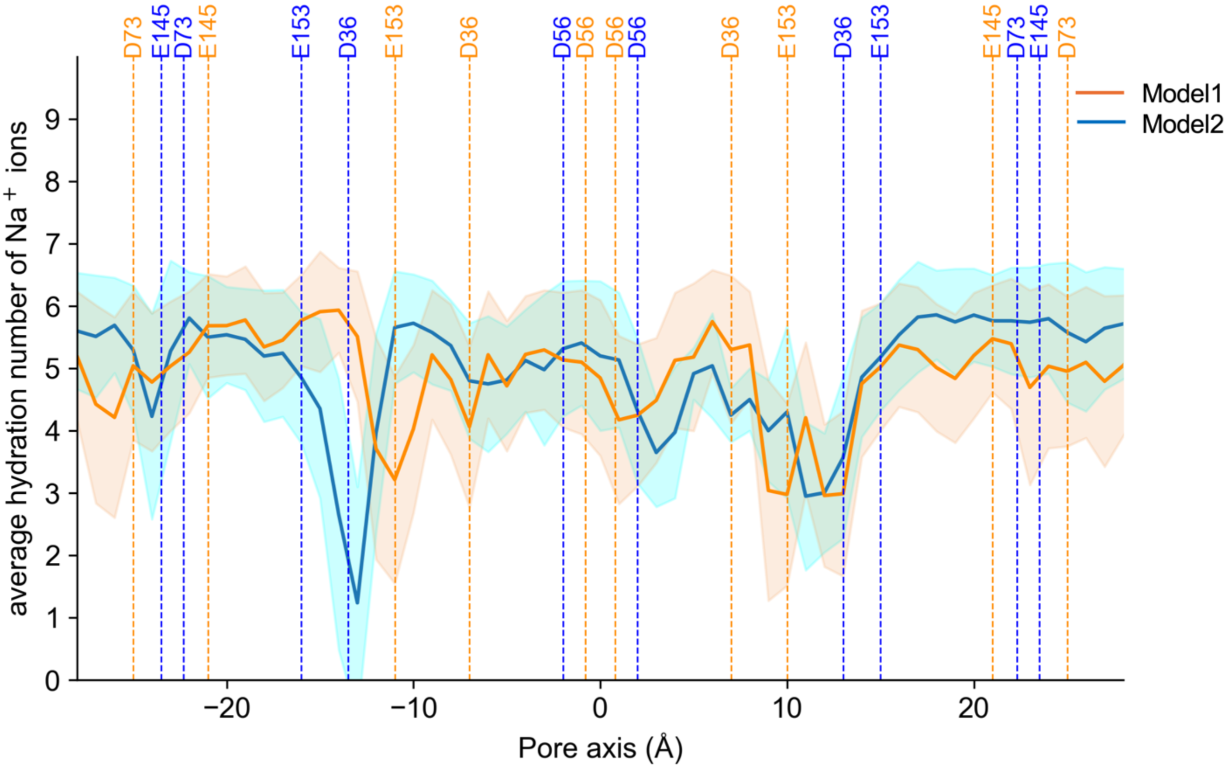


***Figure S4. Average hydration of Na$^+$ ions inside the Claudin-10b paracellular cavity.*** *The average number of Na$^+$ coordinating water molecules along the pore axis is shown in orange for Model1 and blue for Model2. The profile and the associated error are calculated as mean and standard deviation over the three replicas of each model. The positions of the acidic pore-lining residues are referred to those of the corresponding Cα atoms in the equilibrated structure.*

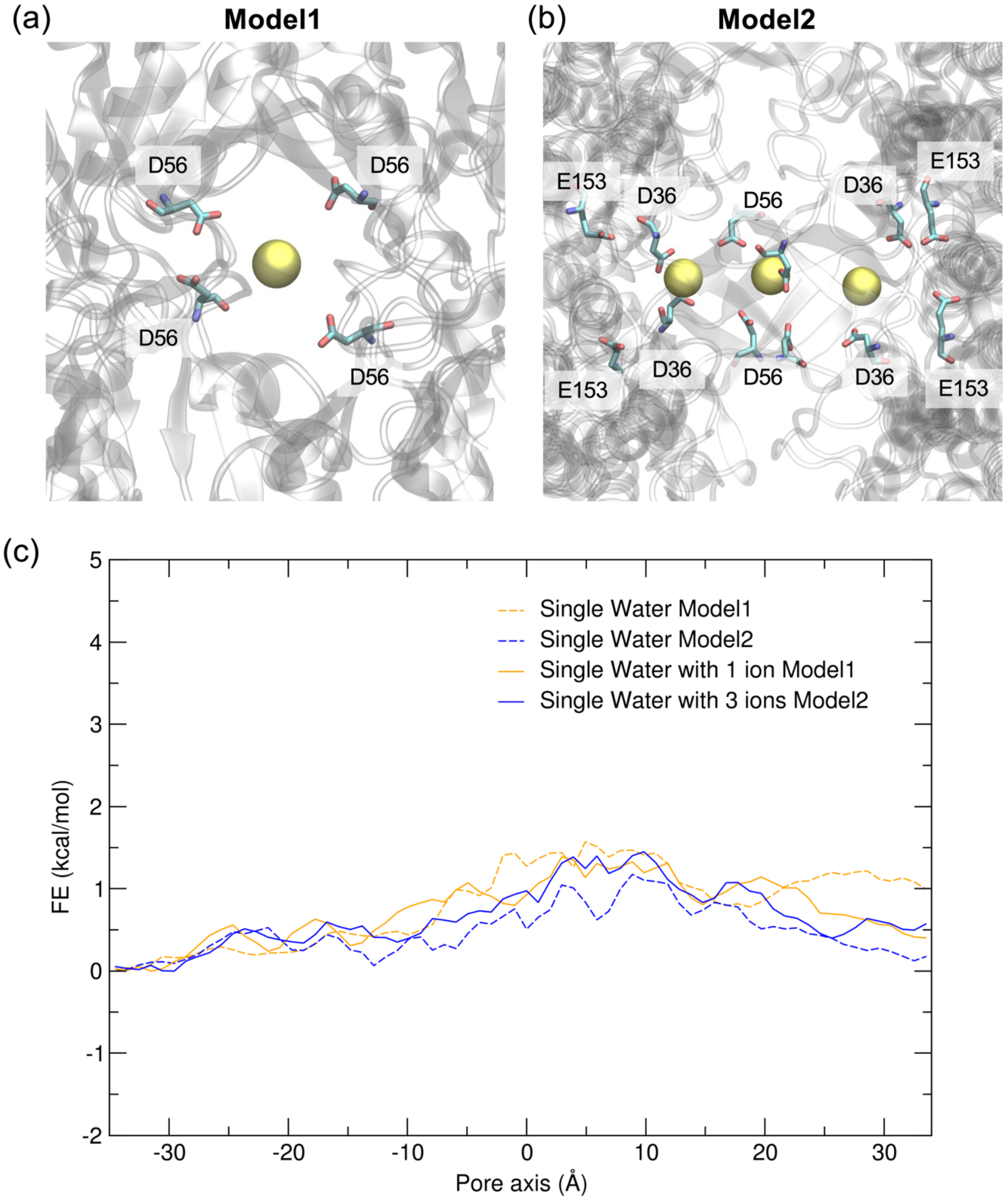


***Figure S5. Free energy calculations of water permeation with $Na^+$ ions coordinated to the acidic sites.*** *(a) Representative snapshot of the single $Na^+$ coordinated to the D56 ring during the US simulations of water permeation through Model1. (b) Snapshot of the three $Na^+$ ions coordinated to the D56 ring and the adjacent D36 pairs during the US simulations of water permeation through Model2. (c) FE profiles of water permeation through partially occluded cavities are reported as solid lines colored in orange for Model1 and blue for Model2, and compared to the results for single water molecule permeations (dotted lines).*

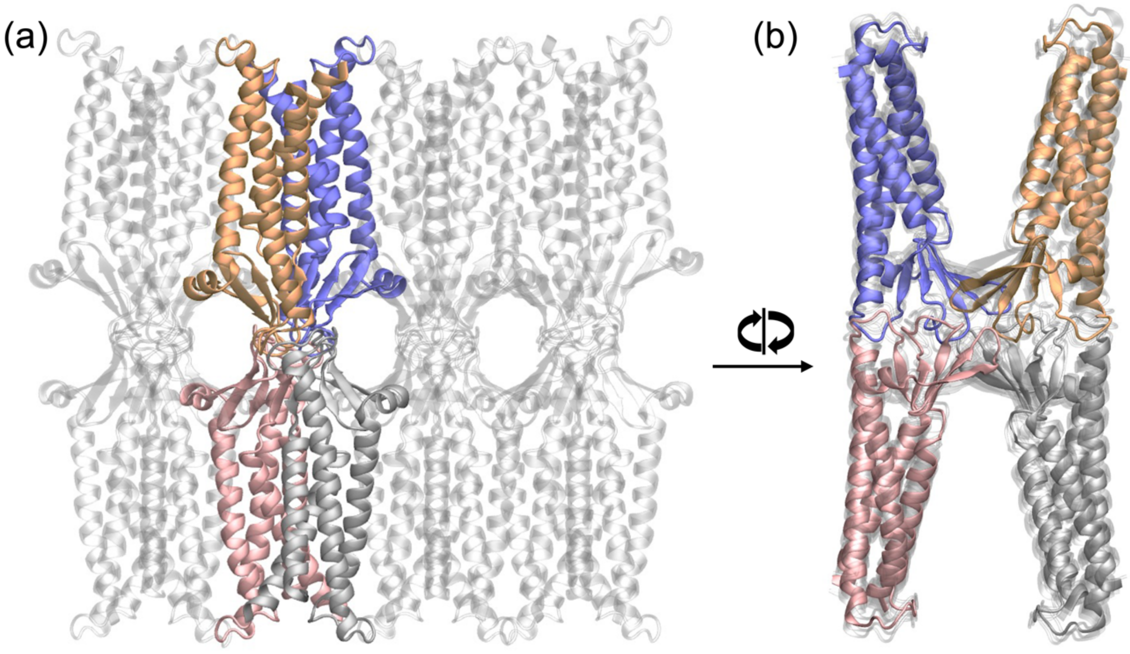


***Figure S6. Minimal unit of the triple-pore models.*** *(a) Frontal and (b) lateral views of Model1, in which it is highlighted the unit forming the interface between two adjacent pores. This structural template was selected from the Cldn15 double-pore model [6] and used to reproduce the Cldn10b triple-pore.*

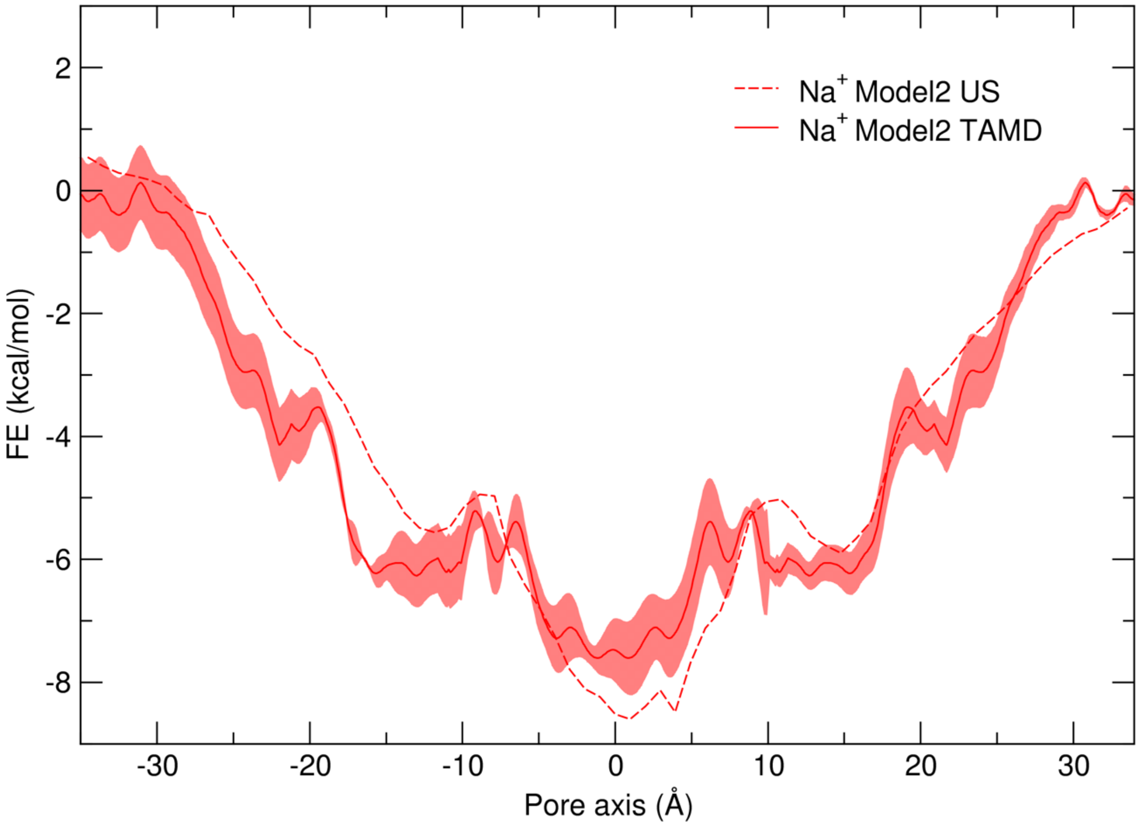


***Figure S7. Comparison between FE profiles of $Na^+$ permeation performed with TAMD and US.*** *The FEs of single $Na^+$ ion permeation through the Model2 central cavity are shown as red solid line for TAMD calculations and as red dotted line for US-WHAM. The FE curve and associated errors are reported as mean and standard deviation over two replicas.*

**Table S1.** Simulation time, timestep and restraints applied during each phase of the equilibration procedure.

| Equilibration Phase | Simulation time (ns) | Time step (fs) | Restrained atoms | Harmonic constant (kcal/ (mol · Å$^2$)) |
|---|---|---|---|---|
| **Phase 1** | 10 | 1 | Heavy atoms of the protein. | 10 |
| **Phase 2** | 10 | 1 | Backbone of the TM helices and the entire ECL domain. | 10 |
| **Phase 3** | 10 | 2 | Backbone of the TM helices and the entire ECL domain. | 10 |
| **Phase 4** | 10 | 2 | C$\alpha$ atoms of the TM amino acids 7, 10, 13, 17, 20, 23, 75, 78, 81, 93, 96, 99, 138, 141, 144, 111, 114, 117, 161, 164, 167, 174, 177, 181 and the entire ECL domain. | 10 |
| **Phase 5** | 10 | 2 | C$\alpha$ atoms of the TM amino acids 7, 10, 13, 17, 20, 23, 75, 78, 81, 93, 96, 99, 138, 141, 144, 111, 114, 117, 161, 164, 167, 174, 177, 181, the unfolded ECL segments and the backbone of the $\beta$-sheet domain. | 10 |
| **Phase 6** | 10 | 2 | C$\alpha$ atoms of the TM amino acids 7, 10, 13, 17, 20, 23, 75, 78, 81, 93, 96, 99, 138, 141, 144, 111, 114, 117, 161, 164, 167, 174, 177, 181 and of the $\beta$-sheet domain, and the unfolded ECL segments. | 2 |
| **Phase 7** | 10 | 2 | C$\alpha$ atoms of the TM amino acids 7, 10, 13, 17, 20, 23, 75, 78, 81, 93, 96, 99, 138, 141, 144, 111, 114, 117, 161, 164, 167, 174, 177, 181 and the unfolded ECL segments. | 2 |
| **Phase 8** | 10 | 2 | C$\alpha$ atoms of the TM amino acids 7, 10, 13, 17, 20, 23, 75, 78, 81, 93, 96, 99, 138, 141, 144, 111, 114, 117, 161, 164, 167, 174, 177, 181 and of the unfolded ECL segments. | 2 |
| **Phase 9** | 10 | 2 | C$\alpha$ atoms of the TM amino acids 7, 10, 13, 17, 20, 23, 75, 78, 81, 93, 96, 99, 138, 141, 144, 111, 114, 117, 161, 164, 167, 174, 177, 181. | 2 |
| **Phase 10** | 10 | 2 | C$\alpha$ atoms of the TM amino acids 7, 10, 13, 17, 20, 23, 75, 78, 81, 93, 96, 99, 138, 141, 144, 111, 114, 117, 161, 164, 167, 174, 177, 181. | 2 |
